# An Integrated Population Model to Incorporate Spatio-Temporal Heterogeneity in Demographic Rates

**DOI:** 10.64898/2026.03.03.709263

**Authors:** Fabian R. Ketwaroo, Matia H. Muller, James F. Saracco, Michael Schaub

**Affiliations:** Swiss Ornithological Institute, Sempach, Switzerland; The Institute for Bird Populations, Petaluma, California, USA

**Keywords:** Change of support, Data integration, Demographic Rates, Nearest Neighbor Gaussian Process, Spatially Explicit, Data Misalignment

## Abstract

1. Demographic processes in populations are inherently heterogeneous across both space and time. Many ecological models explicitly account for temporal heterogeneity in the demographic rates that govern these processes, but assume spatial homogeneity. Ignoring spatial heterogeneity can bias inference, limit predictive performance, and obscure key spatial structure in demographic rates. Integrated population models (IPMs) offer a powerful framework to estimate spatio-temporal demographic rates by combining diverse ecological data sources collected from multiple sampling locations. However, to accomplish this, IPMs face significant statistical and computational hurdles, including misalignment between different data sources and the need to efficiently account for residual spatial autocorrelation.
2. We present a novel Bayesian spatially explicit integrated population model (sIPM) which integrates population count and capture–recapture data from multiple sampling locations to estimate and predict continuous spatio-temporal demographic rates, such as survival, recruitment and population growth rate, across large geographic domains. This framework employs a joint likelihood approach with change of support to flexibly accommodate spatial and spatio-temporal data misalignment, and incorporates a nearest-neighbor Gaussian process to efficiently model residual spatial autocorrelation and generate spatial predictions.
3. We assess the performance of our sIPM through an extensive simulation study. Results show that our approach provides unbiased and precise estimates and predictions of spatio-temporal demographic rates, even in the presence of significant data misalignment and residual spatial autocorrelation. We demonstrate the utility of our method by analyzing data on Gray Catbirds (*Dumetella carolinensis*) from the North American Breeding Bird Survey and the Monitoring Avian Productivity and Survivorship program across the eastern coast of the United States from 2004-2014. This analysis results in maps of apparent survival, recruitment and population growth rate, thereby revealing important spatio-temporal variations in demographic rates that would have been obscured by traditional, spatially homogeneous IPMs.
4. Our sIPM offers a robust and computationally efficient method for studying spatio-temporal variation in demographic processes across large areas, even in the presence of data misalignment and residual spatial autocorrelation. Ultimately, this framework, applicable to many ecological monitoring programs, facilitates the development of spatially targeted strategies necessary for effective conservation and management.

## 1 Introduction

Demographic processes such as birth and death vary across space and time in response to heterogeneity in habitat quality, intra-and interspecific competition, or food supply (Coulson et al., 2001; Gaillard et al., 2000). Quantifying this spatio-temporal variation in demographic processes is a central challenge in ecology and conservation. Nevertheless, ecological models often assume that demographic rates (e.g., survival, recruitment) that govern these processes vary over time but are spatially homogeneous. Ignoring spatial heterogeneity in demographic rates can result in biased inferences, reduce predictive accuracy and obscure understanding of how demographic rates vary spatially (Holt, 1984).

Integrated population models (IPMs; Besbeas et al. (2002), Fournier and Archibald (1982), Schaub and Kéry (2022)) are a popular class of statistical models that are well suited to provide a comprehensive framework to estimate spatio-temporal demographic rates. IPMs integrate multiple ecological data sources (hereafter, data sources), such as individual-level data (e.g. capture-recapture data) and population-level data (e.g. population count data) within a single robust statistical framework. This allows demographic processes to be studied by simultaneously estimating demographic rates and population abundance. In comparison to statistical models that analyze each data source independently, IPMs take advantage of the information shared between data sources to improve parameter estimation and enable the estimation of parameters for which no direct data are available (e.g., immigration (Abadi et al., 2010)).

To date most IPMs are not spatially explicit and assume that demographic rates are temporally but not spatially heterogeneous. However, with the increase in the availability of different spatio-temporally replicated data across large spatial domains in biodiversity monitoring programs (Zipkin et al., 2021) and advancements in computation power and spatial statistics, there is motivation to develop of spatially explicit IPMs that account for the geographic locations of individual populations to estimate and predict spatio-temporal demographic rates. Even so, to accomplish this, there are a number of statistical and computational challenges that need to be addressed.

Data sources often differ in their spatial support (i.e., the spatial domain (point or area) over which an observation or set of observations is collected or aggregated), measurement error, sampling design, and sampling bias (Dungan et al., 2002). For example, demographic studies, such as capture-recapture, are typically conducted at a small spatial support and at a limited number of sampling locations (hereafter, locations) due to the high cost and effort involved, whereas studies such as counting individuals are often conducted at a larger spatial support and across a greater number of locations. Such differences in data sources can lead to spatial data misalignment, meaning that data sources are observed over different spatial supports and/or at non-coincident locations. Additionally, when sampling does not occur simultaneously across data sources, for instance when new locations are added or existing ones are discontinued, data can also become temporally misaligned. In practice, data sources are usually spatiotemporally misaligned, making accurate integration of data sources within IPMs essential (Frost et al., 2023). Misspecification in data integration can propagate biases and uncertainty through the model, leading to incorrect inferences and unreliable predictions. Notably, integrating temporally misaligned data sources within IPMs is well established. In contrast, integrating spatially misaligned data sources within IPMs remains poorly understood and addressed (Pacifici et al., 2019). Ignoring spatial data misalignment by, for example, aggregating all data sources to a common larger spatial support can obscure fine-scale spatial variation, bias parameter estimates, reduce predictive power and inflate uncertainty in demographic rates. This loss of information is well known in spatial statistics (A. E. Gelfand et al., 2001). Addressing spatio-temporal data misalignment in IPMs allows all available data to be used at their native spatial support, broadens the spatio-temporal scope of inference and improves parameter estimation and predictive performance.

Another key challenge is accounting for the spatial autocorrelation of demographic rates that cannot be explained by available covariates. According to Tobler’s first law of geography (Tobler, 1970), demographic processes of individuals that are nearby in space and time tend to be more similar than those far apart. Thus, demographic rates can be spatially correlated due to the complex, unobservable processes that drive them. Failing to account for such residual (unexplained) spatial autocorrelation can result in biased estimates of parameter precision and reduce predictive power (Haining & Li, 2020). Spatially explicit random effects can be used to capture residual spatial autocorrelation. However, this approach increases model complexity and presents computational challenges. This is especially true when modeling locations in continuous space (i.e in point-referenced spatial regression models) as computational complexity increases cubically with the number of locations (the ‘big N’ problem (Banerjee & Fuentes, 2012)).

More recently, attempts have been made to develop spatially explicit IPMs, primarily in the work of Zhao (2020) and Prochazka et al. (2024). While these studies produce spatio-temporal estimates of demographic rates, they do not account for spatial data misalignment or residual spatial autocorrelation. Zhao (2020) aggregated data across large geographic domains to alleviate spatial data misalignment. Although this method is computationally efficient, it assumes homogeneity within the aggregated areas, despite the possible existence of heterogeneous demographic processes. Furthermore, Bosley et al. (2022) showed that such aggregation can lead to estimation bias when demographic processes are spatially heterogeneous. Prochazka et al. (2024) grouped locations that were in agreement with each demographic study considered within constructed minimum convex polygons (MCPs). They then used Bayesian inverse distance weighting (IDW; Burrough et al. (2015)) to predict missing demographic rates at locations where data was lacking, thus increasing the spatial coverage of inference. While this approach is innovative, it also assumes homogeneity within MCPs. Additionally, Prochazka et al. (2024) highlighted that predictions are sensitive to the level of spatial autocorrelation between locations. This is due to the prediction method employed. IDW does not consider the spatial autocorrelation structure, it only considers distance. Notably, both studies relied solely on spatial covariates to model demographic rates. However, the covariates used are unlikely to fully explain the spatial variation in demographic rates, which may lead to biased parameter estimates and reduced predictive accuracy.

Here, we develop a Bayesian spatially explicit integrated population model (sIPM) designed to overcome these challenges, enabling robust estimation at native spatial supports and high resolution prediction of spatio-temporal demographic rates across large geographic domains. Through the power of a joint likelihood formulation with change of support (Cressie, 2015), our framework integrates two popular data sources: population count data and capture-recapture (CR) data in a flexible way to handle spatial and spatio-temporal data misalignment. Population count and CR data provide complementary insights into population dynamics. To account for residual spatial autocorrelation, we use a nearest-neighbor Gaussian process (NNGP; Datta, Banerjee, Finley, and Gelfand (2016)), a computationally efficient approach that approximates a Gaussian process (GP; Williams and Rasmussen (2006)) while drastically reducing computation times (Finley et al., 2019) without sacrificing its key benefits. GPs are widely considered as the gold standard for spatially explicit random effects due to their flexible non-parametric nature. They not only capture complex spatial dependencies but also provides probabilistic uncertainty quantification and robust predictive inference. Compared to simpler methods such as IDW, NNGP allow efficient predictions via kriging (Cressie, 1990), a geostatistical method that accounts for both distance and the underlying spatial autocorrelation structure. Consequently, this framework can efficiently estimate and predict continuous spatio-temporal demographic rates such as survival, recruitment and population growth rate.

We conduct an extensive simulation study to assess the estimation accuracy and predictive capabilities of the proposed modeling framework in the presence of various degrees of data misalignment and residual spatial autocorrelation. Finally, we demonstrate the utility of our sIPM by jointly analyzing count data from the North American Breeding Bird Survey (BBS) and CR data from the Monitoring Avian Productivity and Survivorship program (MAPS), two complementary large-scale monitoring programs. Focusing on the Gray Catbird (*Dumetella carolinensis*) along the eastern coast of the United States from 2004 to 2014, our framework provides important insights into the spatio-temporal heterogeneity in demographic rates and relative population abundance.

## 2 Materials/Data

We integrate two complementary demographic data sources: count data and capture-recapture (CR) data collected at multiple locations and sampling occasions (hereafter, occasions). Count data provide observations of the number of individuals observed at each location and occasion, offering information about population abundance and population trends. CR data consist of individual encounter histories from marked or uniquely identified individuals collected at locations across the study period, providing information on demographic rates. Locations can be points, areas, transects, etc., and occasions can be years, months, etc. Within a IPM, information from these two sources of data are formally shared to improve the estimation of demographic rates and population abundance (Schaub & Kéry, 2022).

## 3 Methods

### 3.1 Demographic models

We focus on the life cycle of a species that starts to reproduce at an age of 1 year. The IPM is composed of a state-space model for the count data and a Cormack-Jolly-Seber (CJS) model for the CR data (Lebreton et al., 1992).

#### 3.1.1 State space model

For *i* = 1, …, *M*_*C*_ count locations, *t* = 1, …, *T* − 1 occasions, we define the state process as follows

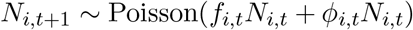

where *ϕ*_*i,t*_ is apparent survival (i.e., the probability that an individual survives and remains at the same location; hereafter, survival) at location *i* from occasion *t* to *t*+1 and *f*_*i,t*_ is the fraction of new individuals (recruits) at location *i* and occasion *t* + 1 relative to the population abundance (*N*_*i,t*_) at location *i* and occasion *t*. Recruitment is a function of productivity, juvenile survival, site fidelity, and immigration. The population growth rate at location *i* and time *t* + 1 is derived and calculated as the ratio of *N*_*i,t*+1_ to *N*_*i,t*_, or as the sum of recruitment and survival. Hereafter, we refer to population abundance and population growth rate as abundance and growth rate, respectively. For each location *i*, the abundance at occasion 1 (*N*_*i*,1_) is latent. For *t* = 1, …, *T* occasions, the observation process is defined as

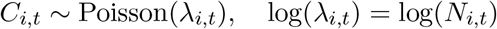

where *C*_*i,t*_ is the observed counts at location *i* occasion *t* and *λ*_*i,t*_ is the expected number of counts at location *i* occasion *t*. Naturally, *λ*_*i,t*_ is a function of *N*_*i,t*_ and can also include covariates to account for observation error. Probability distributions that account for overdispersion such as the Negative Binomial distribution can also be used.

#### 3.1.2 Cormack Jolly Seber (CJS) Model

For computational reasons, we employ the m-array formulation of the CJS model (Schaub & Kéry, 2022). For *i* = 1, …, *M*_*CR*_ CR locations, the likelihood is expressed as

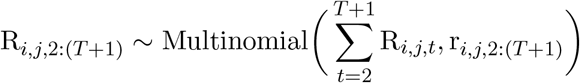

where R_*i,j,t*_ denotes the number of individuals released at location *i* occasion *j* that are next recaptured at location *i* occasion *t* (*j* = 1, …, *T* − 1, *t* = 2, …, *T, t* > *j*), and R_*i,j,T* +1_ denotes the number of individuals released at location *i* occasion *j* and never recaptured again at location *i*. The cell probabilities r_*i,j*,2:(*T* +1)_ represent the expected frequency of each capture history fragment at location *i* which is expressed as a function of survival (*ϕ*_*i,t*_) at location *i* from occasion *t* to *t* + 1 and recapture probability (*p*_*i,t*_) at location *i*, occasion *t* + 1:

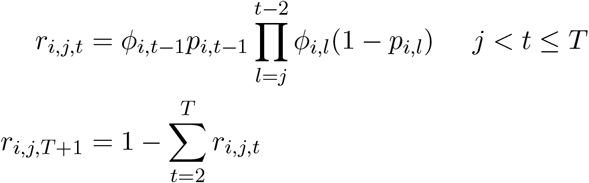

### 3.2 Data integration

Data integration refers to the process of combining information from various sources within a single statistical modeling framework to create a comprehensive understanding of complex and interacting ecological processes (Michener & Jones, 2012; Schaub & Abadi, 2011) such as population dynamics, species distribution, etc. It assumes that each data source provides information, directly or indirectly, about one or more components of the ecological process of interest. By exploiting this information, data integration can overcome limitations of individual datasets analysis, i.e., estimate confounded parameters (Cole & McCrea, 2016), improve parameter precision and accuracy (Schaub & Abadi, 2011) and increase spatio-temporal coverage of inference (Saunders et al., 2019).

In IPMs, a joint likelihood approach is used to combine multiple data sources into a unified statistical framework. Let ***y***_*i*_, *i* = 1, …, *n* distinct population data sets associated with a model ℳ_*i*_ with parameter vector ***θ***_*i*_. Assuming conditional independence of the data sources given its model parameters, the joint likelihood is defined as

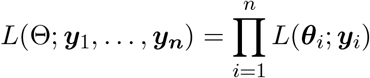

where *L*(***θ***_*i*_; ***y***_*i*_) denotes the likelihood of ***y***_*i*_ and 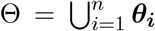 is the full parameter set. It is further assumed that some components of ***θ***_***i***_ are the same in multiple data sources (e.g. demographic rates), such that ***θ***_***i***_ ∩ ***θ***_***j***_ ≠ ∅ for *i* ≠ *j*, allowing information to be shared across data sources. Specification of *L*(***θ***_*i*_; ***y***_*i*_) reflect the relationships between different data sources and the underlying parameters in the ecological process model. Based on this joint likelihood formulation, we illustrate how to integrate counts and CR data under three data integration scenarios: perfect data alignment, spatial data misalignment, and spatio-temporal data misalignment.

#### 3.2.1 Perfect data alignment

Perfect data alignment refers to the case where there are equal types of data sources at all locations and occasions, rendering the number of count and CR locations and occasions to be the same. For *i* = 1, …, *M* locations, *t* = 1, …, *T* occasions, the joint likelihood is expressed as

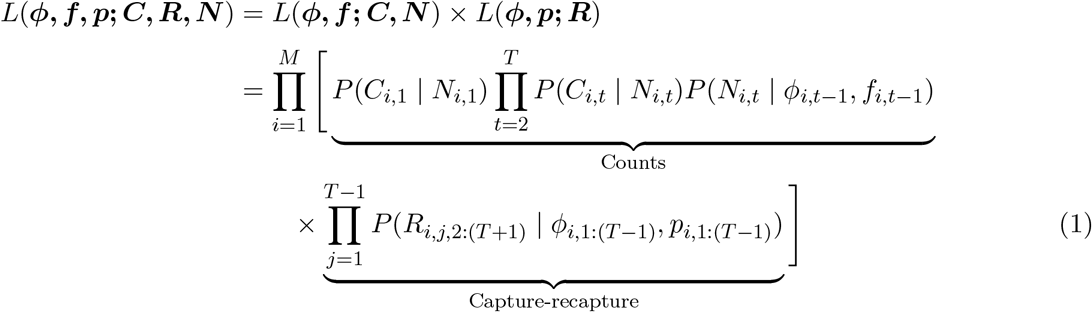

where *P* (·|·) denotes the probability function of the data given its latent parameters. From equation 1, we can see that ***ϕ*** occurs in both data source likelihoods, allowing information to be shared across both data sources. Importantly, this allows ***f*** to be estimated more accurately and avoids estimation difficult present in models that only use count data (insufficient information in the data (Hostetler & Chandler, 2015)).

#### 3.2.2 Spatial data misalignment

In a spatial context, data sources are often observed at non-coincident locations and over different spatial supports. To integrate spatially misaligned data, we employ the concept of change of support (COS; Cressie (2015) and Gotway and Young (2002)) from spatial statistics. COS reconciles spatially misaligned data by linking each data source to an underlying shared spatial stochastic process through its native spatial support.

A spatial stochastic process is a collection of random variables defined over a spatial domain (Pavliotis, 2014). Let *Z*(·) ≡ {*Z*(*s*) : *s* ∈ *D*} denote a spatial stochastic process defined on a continuous spatial domain *D* ⊂ ℝ^*d*^. Under COS, this process is expressed over a different spatial support *B* ⊂ *D*, where *B* represents any region or geometry included in *D*. This results in another process {*Z*(*B*)}, where *Z*(*B*) represents an aggregate (typically an average or total) of *Z*(·) over *B*. By definition

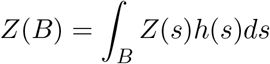

where *h*(*s*) is a weighting function that determines the importance of each location *s* within *B*. This weighting function is often defined as |*B*|^−1^ to obtain an average over *B*, or 1 for a total. Here we assume the latter. In this way, each data source is linked to the continuous process *Z*(·) through its own spatial support *B*, which defines *Z*(*B*). This formulation allows spatially misaligned data sources to be integrated without aggregation to a common spatial support. Thus, it is important to use stochastic processes *Z*(·) that allow for efficient and accurate computation of *Z*(*B*). Therefore, we assume *Z*(*s*) = ***X***(*s*)^⊤^***β*** + *ω*(*s*) where ***X***(*s*) is a vector of spatial covariates including the intercept, ***β*** is the corresponding vector of regression coefficients and *ω*(*s*) is the residual spatial process. For clarity, assume two spatially misaligned datasets *Y*_*b*_ = {*Y* (*b*_*i*_) : *i* = 1, …, *n*_*b*_ locations} and *Y*_*c*_ = {*Y* (*c*_*i*_) : *i* = 1, …, *n*_*c*_ locations}. To integrate spatially misaligned data through COS, it is assumed that a *Z*(·) exists and that each dataset is a realization of this underlying process over its respective spatial support *B*. Conditional on *Z*(·), the joint likelihood is expressed as

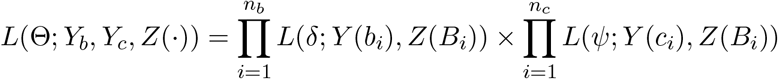

where Θ = (*δ, ψ*) is the parameter set. This formulation resolves spatial data misalignment and makes use of the strengths of each dataset, providing a more robust and comprehensive understanding of the underlying spatial process, particularly in areas where data are sparse. Similarly, spatially misaligned data with temporal replication can be combined assuming there is a spatio-temporal stochastic process connecting data sources. Let *Z*(·, ·) ≡ {*Z*(*s, t*) : *s* ∈ *D, t* = 1, …, *T*} be a spatio-temporal stochastic process. To account for spatial misalignment of temporally replicated count and CR data in our sIPM, we let

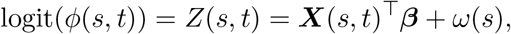

where ***X***(*s, t*) is a vector of spatio-temporal covariates including the intercept, such that at each location *s*_*i*_: *ϕ*_*i,t*_ := *ϕ*(*s*_*i*_, *t*) = logit^−1^ *Z*(*s*_*i*_, *t*) for *i* = 1, …, (*M*_*C*_ + *M*_*CR*_) locations, *t* = 1, …, *T* − 1 occasions. Consequently, the joint likelihood is written as

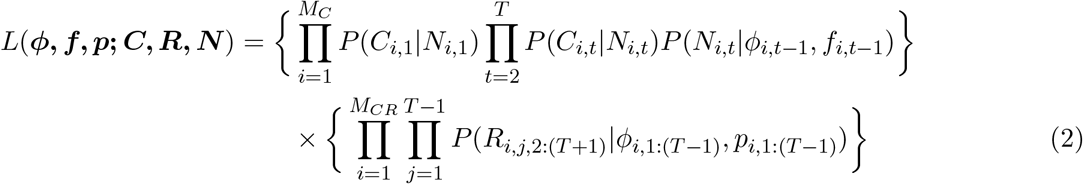

This formulation uses the detailed information about ***ϕ*** provided by the CR data via the CJS model to help inform and constrain ***ϕ*** estimates in locations where only count data are available. It also improves the estimation of ***f*** as stronger information on ***ϕ*** helps to distinguish between the two demographic rates. Consequently, this formulation expands the spatio-temporal scope of inference in a robust manner.

#### 3.2.3 Spatio-temporal data misalignment

Spatio-temporal data misalignment occurs when spatially misaligned data are inconsistently sampled across all occasions. In this case, we adopt the COS framework used to integrate spatially misaligned data and extend it to accommodate spatio-temporal misalignment with minor refinements. As such, the joint likelihood is expressed as

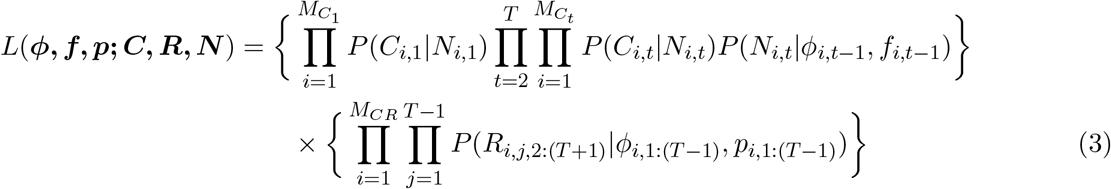

where 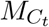 denotes the number of count locations sampled at occasion *t*. For CR locations, we set the *p*_*i,t*_ = 0 for location-occasion combinations where no sampling was conducted.

### 3.3 Demographic rates

In this section, we describe how demographic rates are modeled in the three data integration scenarios. Assuming perfect data alignment, for *i*, …, *M* locations, *t* = 1, …, *T* − 1 occasions we model survival and recruitment as:

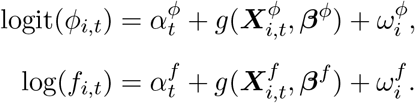

For notational convenience, let *ν* ∈ (*ϕ, f*), such that 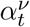 denotes an occasion-specific baseline (intercept on the link scale) that is independent across occasions, 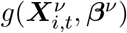 is a generic function that relates covariates 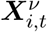 (*ℓ*^*ν*^ dimensional vector) to a corresponding vector of regression coefficients ***β***^*ν*^ and 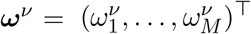 is a spatial random effect vector that accounts for spatial residual autocorrelation. Under spatial data misalignment, *M* = *M*_*C*_ + *M*_*CR*_ and under spatio-temporal data misalignment, the number of locations at occasion *t* is 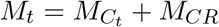. Importantly, given data misalignment, we only estimate recruitment at count locations while survival is estimated at both the count and the CR data locations. Table 1 summarizes the locations and parameter estimability under each scenario.

**Table 1:** Summary of the number of locations available and of the number of locations from where survival and recruitment is estimated under different data-integration scenarios. *M* = total number of locations available, *M*_*C*_ = number of count locations, *M*_*CR*_ = capture–recapture (CR) locations, *M*_*t*_ = total number of locations available at *t* and 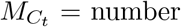 of count locations at time *t*.

| Scenario | Locations available | Survival estimated at | Recruitment estimated at |
| --- | --- | --- | --- |
| Perfect alignment | $M$ | $M$ | $M$ |
| Spatial misalignment | $M = M_C + M_{CR}$ | $M_C + M_{CR}$ | $M_C$ only |
| Spatio-temporal misalignment | $M_t = M_{C_t} + M_{CR}$ | $M_{C_t} + M_{CR}$ | $M_{C_t}$ only |

### 3.4 Gaussian Process (GP)

We model the spatial random effect vector ***ω***^*ν*^ using Gaussian processes (GPs) (Williams & Rasmussen, 2006). A GP is a non-parametric probabilistic modeling framework that defines a probability distribution over possible functions. For simplicity, we omit the subscript *ν* in the following section, as both survival and recruitment are modeled using the same GP specification. Formally, for location *s*,

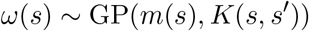

where *m*(*s*) is the mean function and *K*(*s, s*^*′*^) is the covariance function between any two locations (*s* and *s*^*′*^). The mean function gives the expected value of the process at location *s*, while the covariance function specifies the dependence between locations and controls the smoothness and variability of functions drawn from the GP. Critically, for any finite set of spatial locations *H* = {*s*_1_, …, *s*_*M*_}, ***ω*** = (*ω*(*s*_1_), …, *ω*(*s*_*M*_))^⊤^ follows a multivariate normal (MVN) distribution

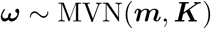

with a mean vector ***m*** = [*m*(*s*_1_), …, *m*(*s*_*M*_)]^⊤^ and covariance matrix ***K*** with entries *K*(*s*_*i*_, *s*_*j*_). This finite representation enables practical computation of GP, as the MVN distribution is well understood mathematically and computationally tractable. Consequently, GPs provide a powerful framework for modeling residual spatial autocorrelation, naturally capturing complex, nonlinear dependence structures in space and yielding reliable parameter estimates with associated uncertainty. In this work, we assume *m*(*s*) = 0 and employ an exponential covariance function:

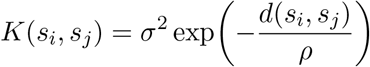

where *d*(*s*_*i*_, *s*_*j*_) is the Euclidean distance between locations *s*_*i*_ and *s*_*j*_, *σ*^2^ is the signal variance parameter that controls the overall amplitude of the function and *ρ* is the length scale parameter that controls how quickly the correlation between locations decays with distance. We use the exponential covariance function because it is simple, provides intuitive parameters, reasonable assumptions about ecological data and is widely used in ecology (Doser et al., 2022, 2024). An added benefit of GPs is that they enable efficient prediction at unobserved locations. The joint distribution of the GP at unobserved locations is also a MVN distribution, yielding both predictions and associated uncertainty (Williams & Rasmussen, 2006).

However, one of the primary challenges of using GPs is their computational complexity and the amount of memory storage required for large datasets of size *M*. In this case, ***K*** is a dense *M* × *M* covariance matrix and model fitting requires the computation of the inverse and determinant of ***K*** which requires 𝒪 (*M*^3^) floating point operations (flops), 𝒪 (*M*^2^) flops for memory storage and predicting the response at *q* new locations require an additional 𝒪 (*qM*^3^) flops. Consequently, GPs can become computationally prohibitive when *M* is even moderately large (e.g., 500). To address this, we employ a sparse approximation method called nearest-neighbor Gaussian process (NNGP; Datta, Banerjee, Finley, and Gelfand (2016)) to model residual spatial autocorrelation in survival and recruitment.

#### 3.4.1 Nearest Neighbor Gaussian Process (NNGP)

The NNGP is a scalable and efficient solution for large spatial data sets by leveraging conditional independence. More formally, the NNGP is a valid GP that is based on expressing ***ω*** as the product of conditional densities

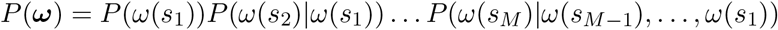

and shrinking the larger conditioning sets on the right-hand side with smaller, carefully chosen, conditioning sets of size *k* ≪ *M*. Consequently for each *s*_*i*_ ∈ *H*, a smaller conditioning set *S*(*s*_*i*_) is used to construct

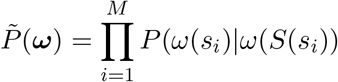

where *S*_*s*_ = {*S*(*s*_*i*_); *i* = 1, …, *M*} is often called the neighbor set. For 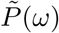 to be a proper MVN distribution, *G* = {*H, S*_*s*_} needs to be a directed acyclic graph (DAG) with *H* as the set of nodes and *S*_*s*_ as the set of directed edges. We accomplish this by following Vecchia (1988) and order *H* along the horizontal axis (longitude or easting) and then select *S*(*s*_*i*_) to be the set of *k* nearest neighbors of *s*_*i*_ with respect to Euclidean distance. Importantly, Datta, Banerjee, Finley, and Gelfand (2016) showed that the effectiveness of the NNGP is largely determined by the number of neighbors specified rather than the specific ordering, particularly for moderate to large neighbor sizes. They demonstrated that *k* = (15, 20) is adequate for most sizes of *M*. In this work, we set *k* = 15. This leads to a valid multivariate normal distribution with a sparse precision matrix 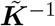 resulting in: 𝒪 (*Mk*^3^) computational cost, 𝒪 (*Mk*^2^) memory storage and 𝒪 (*qk*^3^) for predicting at *q* new locations. The complexity is now linear in terms of *M* and *q*.

### 3.5 Inference

We perform inference in a Bayesian framework using MCMC methods via R package `NIMBLE` (de Valpine et al., 2017) version 1.3.0. For scalable computation with the NNGP, we adopt the efficient algorithms of Finley et al. (2019): pseudocode 2 and 3 for sparsity-inducing computation and evaluation of the quadratic form, and algorithm 2 for posterior predictive inference. To improve computational efficiency, we impose the sum-to-zero constraint on ***ω***^*ν*^, which reduces MCMC autocorrelation among its components. We further used an individual random walk (RW) sampler to sample ***ω***^*ν*^.

## 4 Simulation study

We conducted an extensive simulation study to evaluate the estimation accuracy and predictive power of the proposed modeling framework. Within a unit square study area, we randomly simulated *M* = 500 locations at *T* = 6 occasions, considering three cases of data integration: perfect data alignment (PA), spatial data misalignment (SM) and spatio-temporal data misalignment (STM). For the SM and STM cases, we assume non-coincident locations with 80% of the locations designated as count data and the remaining 20% of locations set as CR data. For the STM case, we considered in addition cases of high and low temporal data misalignment. In the case of low temporal data misalignment, 83.33% and 80% of the occasions were sampled for each count and CR location, respectively. In the case of high temporal data misalignment, 50% and 60% of the occasions were sampled for each count and CR location, respectively. We model demographic rates as described in section 3.3 considering two cases of covariate effects on survival and recruitment: a spatially constant case and a spatially varying case. This allowed us to assess the ability of our modeling framework to recover regression coefficients under both spatially constant and spatially varying covariate effects. For the spatially constant case, we used a linear function with a single spatially correlated covariate: 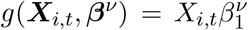, and for the spatially varying case, we employed a interaction function that uses a single spatially correlated covariate interacting with spatial coordinates (latitude (*Lat*) and longitude (*Long*)): 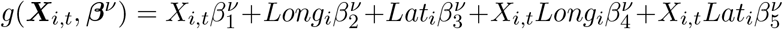. For the PA and SM cases, we only considered the spatially constant covariate effect case, but for the STM case, we consider both the spatially constant and spatially varying covariate effect cases. The spatially correlated covariate and residual spatial autocorrelation were simulated from a GP with exponential covariance function ensuring correlation across large distances. Survival, recruitment and abundance were simulated such that the mean spatial growth rate was 1 with 95% quantile interval of (0.5, 1.5) for each occasion. The recapture probability was modeled as constant across space, but specific to each occasion: 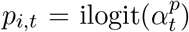 where 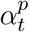 is an independent temporal intercept. Details on the values of the data generating parameter, prior and MCMC settings are reported in section S1.1 of the Supporting Information. Finally, we investigated the predictive performance of the survival and recruitment models in each data integration case at 100 locations for each occasion. Each data integration case was subjected to 100 simulation runs.

To quantify estimation accuracy, we use relative bias (RB), Monte Carlo standard error of mean RB (MCSE(RB)), root mean square error (RMSE) and 95% posterior credible interval coverage (Cov). We further use mean absolute percentage error (MAPE), RB, RMSE and Cov to quantify predictions. See section S1 of Supporting Information for mathematical description of these metrics.

### 4.1 Results

Figure 1 and Table S3 show that demographic rates and abundance are estimated well with low mean RB, low mean RMSE and high mean coverage. As the amount of information decreases due to data misalignment, parameter uncertainty increases but on average remained unbiased. The estimation accuracy is similar regardless of whether the covariate effect was spatially constant or spatially varying. The regression coefficients ***β***^*ν*^ in all cases are estimated well with low mean RB, low RMSE and high coverage (Figure 1 and Tables S4-S5). Additionally, the occasion-specific baseline recapture probability 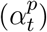, NNGP parameters 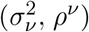, occasion-specific baseline survival and recruitment 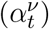 are all estimated well with relatively low RB, low RMSE and high coverage in all cases considered (Tables S4-S5). A notable exception is baseline recapture probability in the terminal occasion 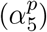. As the level of data misalignment increases, RB increases in 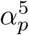 which in turn affects baseline survival estimation in the terminal occasion 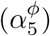 (Figure S1). As the level of data misalignment increases, the strength of information borrowing decreases and our framework reverts to its individual components where it is known that baseline survival and recapture probability are not identifiable in the terminal occasion when using a CJS model with time-dependence in both survival and recapture probability (Lebreton et al., 1992). However, on average, this bias does not significantly impact estimation of spatial survival in the terminal occasion (Table S6).

**Figure 1.**
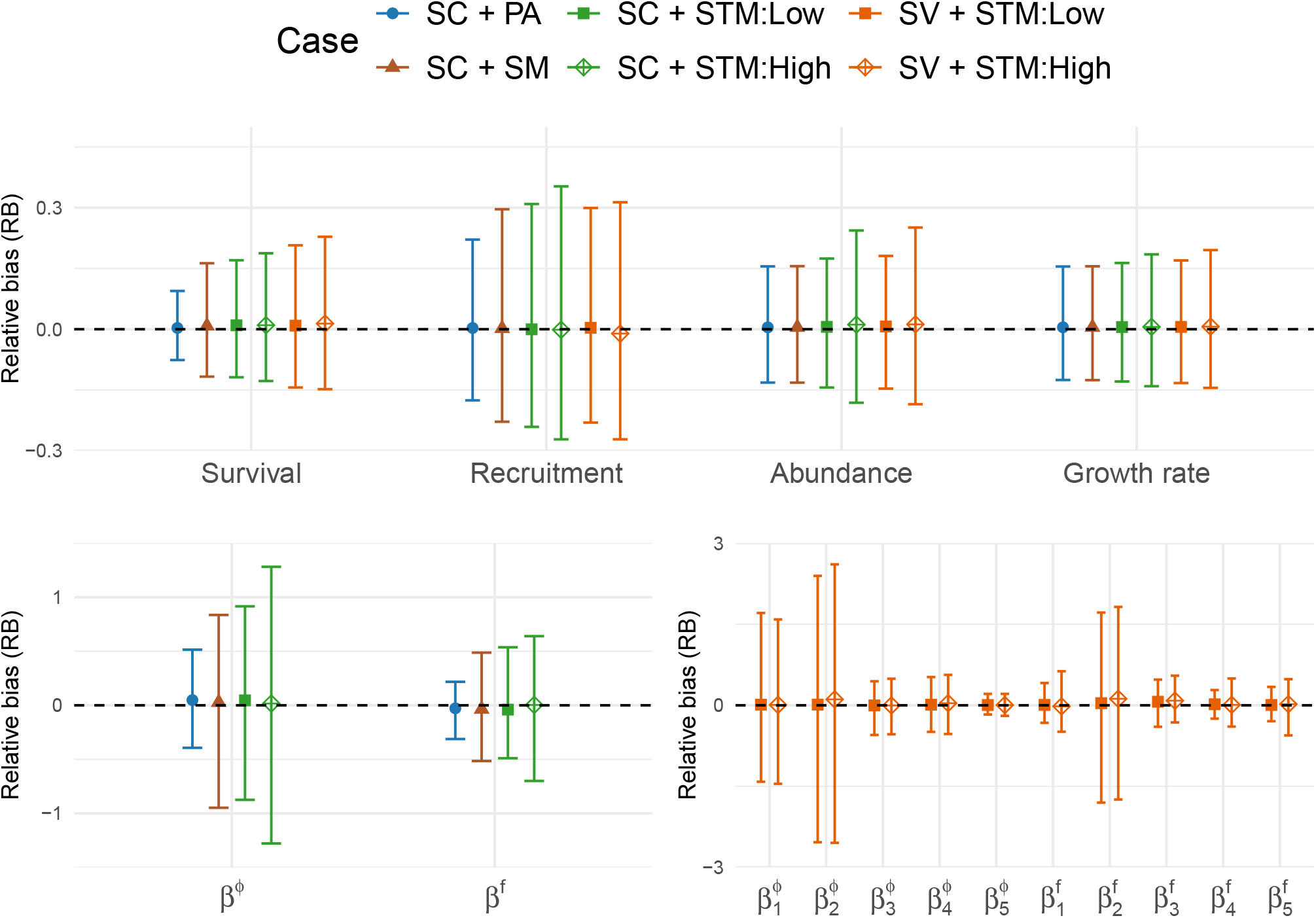
The upper panel displays 95% quantile error bars of mean relative bias (RB) across locations and occasions of survival, recruitment, abundance and growth rate. The lower panels show 95% quantile error bars of RB of regression coefficients for survival (*β*^*ϕ*^) and recruitment (*β*^*f*^). Shapes denote the mean RB across the simulation runs. Results are shown under scenarios with spatially constant (SC) and spatially varying (SV) covariate effects on survival and recruitment, and across data integration cases SC with perfect data alignment (SC + PA), SC with spatial data misalignment (SC + SM), SC with spatial data misalignment with low temporal data misalignment (SC + STM:Low), SC with spatial data misalignment with high temporal data misalignment (SC + STM:High), SV with spatial data misalignment and low temporal data misalignment (SV + STM:Low) and SV with spatial data misalignment and high temporal data misalignment (SV + STM:High).

As shown in Figure 2 and Table S7, survival and recruitment had good predictive power with average MAPE < 20, low RMSE and high mean coverage in all cases. Survival had stronger predictive power than recruitment possibly due to information sharing between datasets. Recruitment with the spatially varying covariate effect seemed to have higher predictive power than recruitment with the spatially constant covariate effect. Similar to estimation accuracy, as the amount of information decreases due to data misalignment, prediction uncertainty increased.

**Figure 2.**
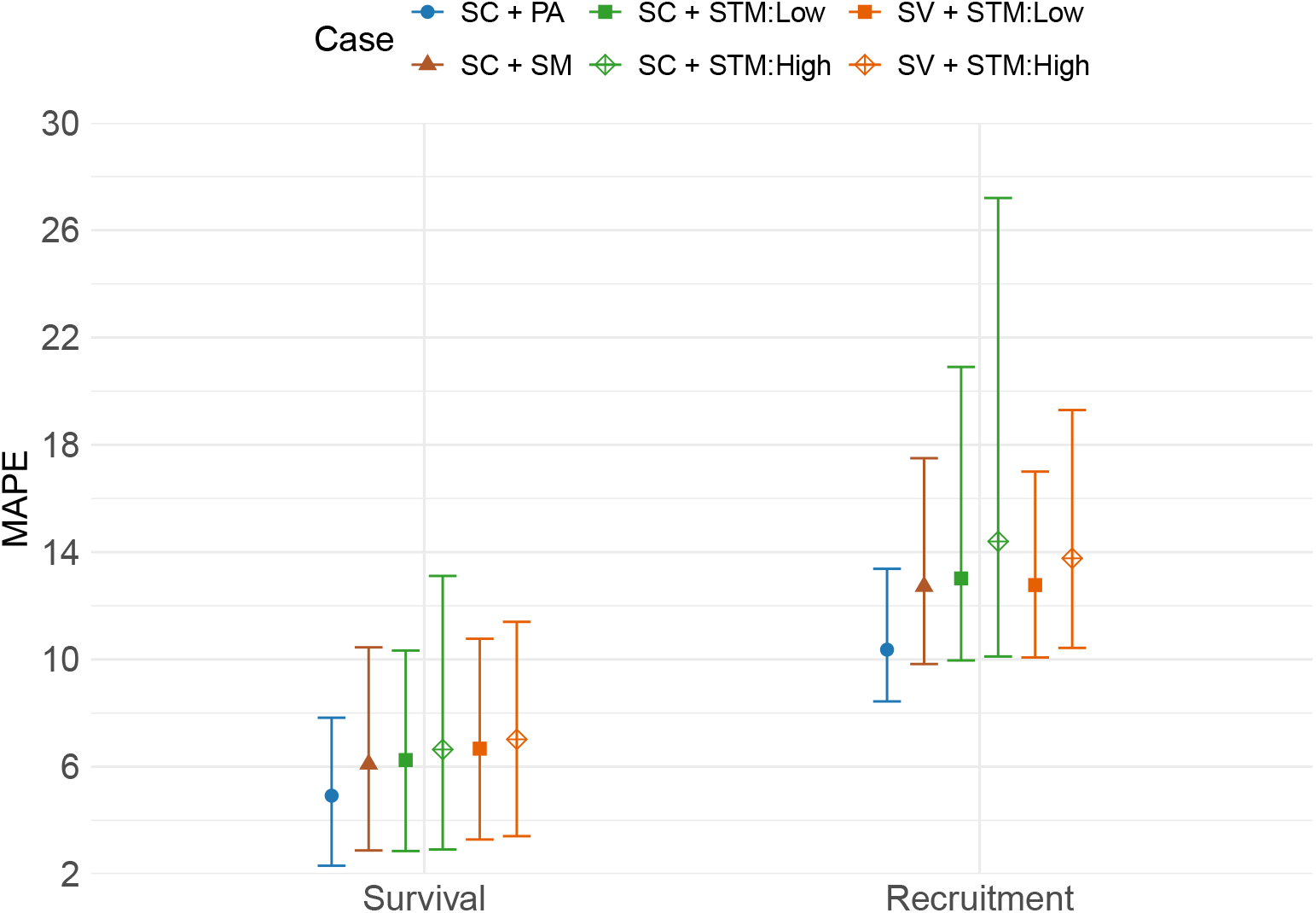
95% quantile error bars of average mean absolute percentage error (MAPE) of survival and recruitment predictions across locations and occasions given spatially constant (SC) and spatially varying (SV) covariate effects under the following cases: SC with perfect data alignment (SC + PA), SC with spatial data misalignment (SC + SM), SC with spatial data misalignment with low temporal data misalignment (SC + STM:Low), SC with spatial data misalignment with high temporal data misalignment (SC + STM:High), SV with spatial data misalignment and low temporal data misalignment (SV + STM:Low) and SV with spatial data misalignment and high temporal data misalignment (SV + STM:High). Shapes denote the mean MAPE across the simulation runs.

## 5 Case study: Gray Catbird

### 5.1 Data

Focusing on the Gray Catbird (*Dumetella carolinensis*), we demonstrate the utility of our sIPM by jointly analyzing count data from the North American Breeding Bird Survey (BBS; David Jr Ziolkowski et al. (2025)) and CR data from the Monitoring Avian Productivity and Survivorship program (MAPS; Desante et al. (1995)). The BBS is a bird monitoring program designed to provide estimates of the relative abundance and population trends of birds in North America. During the breeding season in late May or June each year, breeding birds are counted once at approximately 50 equally spaced points along more than 3,000 randomly established routes, each approximately 39 km long. All birds heard or seen at each point are counted for 3 minutes within a 400 m radius (Sauer et al., 2014). We use the total number of Gray Catbirds detected at all points along a route in a given year as the observed counts for that route and year. MAPS produces CR data by employing a constant effort mist netting protocol at more than 1200 field stations to ring and monitor birds throughout USA and Canada (DeSante & Kaschube, 2009). A MAPS station typically consists of 10 mist nets, each 12 m long, distributed over an area of approximately 8 ha. Stations are operated for up to 9 days per breeding season with each day of operation separated by about 10 days (see DeSante et al. (2004) for additional details).

For consistency, we denote years as occasions, and the centroid of BBS routes and MAPS stations as locations. Accordingly, we considered BBS and MAPS locations along the eastern coast of the United States from 2004 to 2014. More specifically, we included Bird conservation regions 30, 29, 28, 27, 14 and 13. BBS data were obtained using the R package `bbsBayes2` (Edwards et al., 2023) where only locations with non-zero observations were used. MAPS data were obtained using the MAPS Data Exploration Tool (Institute for Bird Populations, 2023), and we excluded locations with fewer than five total captures over the study period. BBS and MAPS data were spatio-temporally misaligned and we only considered locations with more than 4 operation years. Figure 3 shows the spatial distribution of BBS and MAPS locations considered across the time period of interest, 759 locations comprising of 697 BBS locations and 62 MAPS locations.

**Figure 3.**
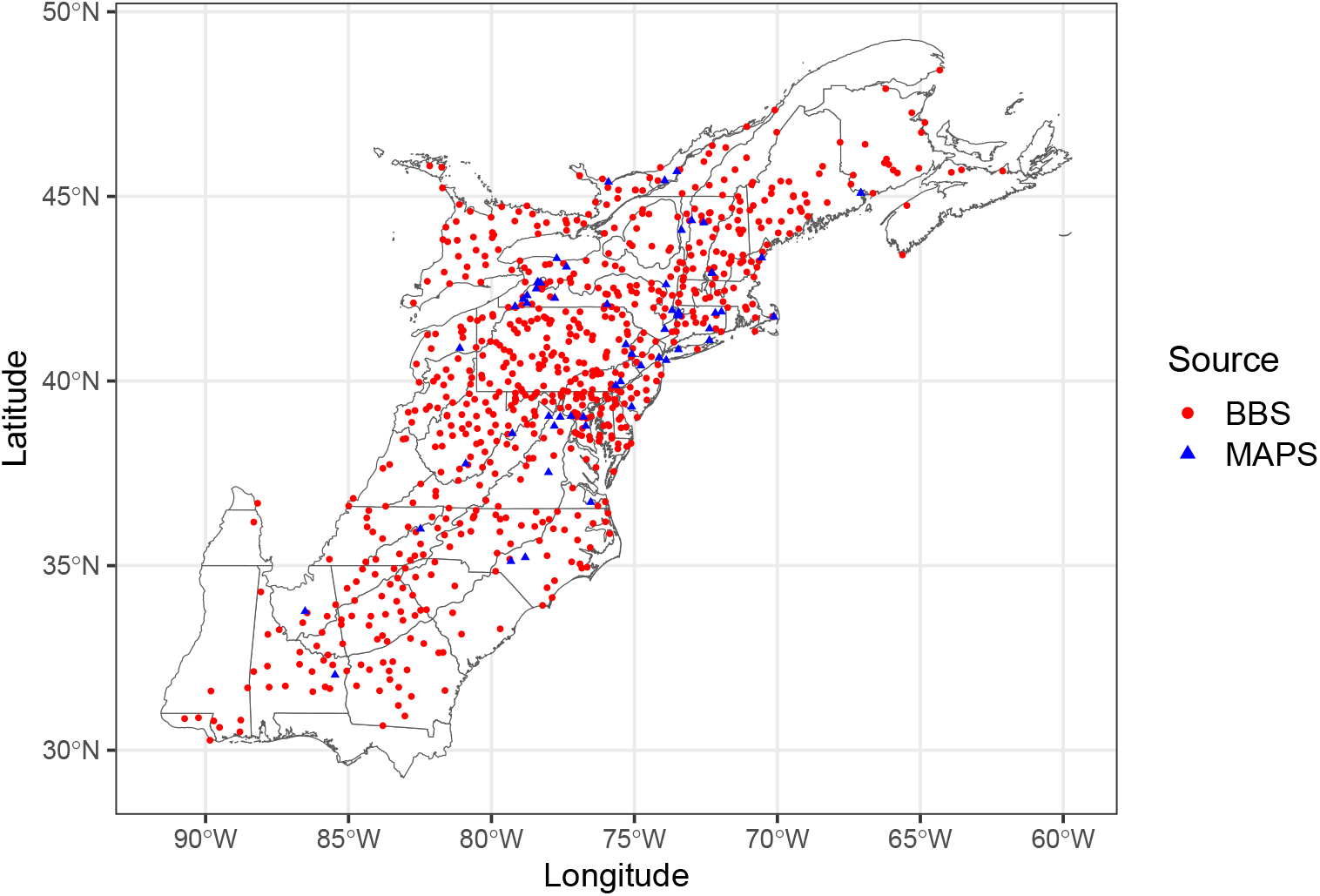
Spatial distribution of BBS (North American Breeding Bird Survey) and MAPS (Monitoring Avian Productivity and Survivorship program) locations in the United States during 2004–2014 as used in the analysis. Circles denote locations of counts from BBS and triangles denote capture-recapture data from MAPS across Bird Conservation Regions 30, 29, 28, 27, 14, and 13 where Grey Catbirds were observed (see section 5.1).

### 5.2 Model specification

For the observation process of the BBS counts, we account for the experience of observers because it is known to affect counts (Link & Sauer, 2002):

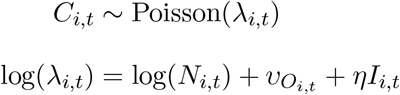

where *C*_*i,t*_ is the observed counts at location *i* on occasion *t*, 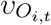 is the observer random effect for observer identity *O*_*i,t*_ at location *i* on occasion *t, I*_*i,t*_ is an indicator variable denoting whether this is the first time an observer is visiting location *i* at occasion *t* and *η* is the first time observer regression coefficient. The observer effect is Gaussian: 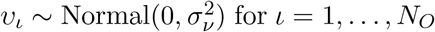 for *ι* = 1, …, *N*_*O*_(number of observers). Importantly, *N*_*i,t*_ is relative abundance at location *i* and occasion *t* as the sampled area of BBS counts are not known. In constant-effort mist surveys such as MAPS, an individual bird that is captured can be either a resident, meaning it belongs to the local breeding population, or a transient, meaning it only uses a location as a temporary stop during migration or settlement. Not accounting for transient individuals results in an underestimation of survival as by definition transient individuals have an survival probability of zero (Pradel et al., 1997). We use a computationally efficient m-array formulation of the CJS model that accounts for transience (Cave et al., 2010; Pradel et al., 1997) to obtain inferences on resident individuals. In this formulation, two m-arrays are used: the first summarizes the capture histories of both residents and transients up to their first recapture occasion (including individuals recaptured within the occasion of the first capture and individuals captured only once), while the second summarizes the histories of residents from the first recapture occasion onward. The second m-array follows the standard m-array CJS model likelihood described in section 3.1.2. The first m-array is an extension of standard CJS model likelihood by incorporating residency probability and probability of confirmation residency status in the cell probabilities to account for transient individuals (see section S2.1 of Supporting Information for mathematical description). We assume Gray Catbirds captured twice during the breading season and at least 7 days apart are known residents. Recapture probability (*p*_*i,t*_) at location *i* and occasion *t* is modeled as

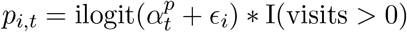

where 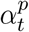 is an independent temporal intercept, *ϵ*_*i*_ is a location random effect and visits is a binary variable denoting if the location was sampled or not, 1 denoting sampled and 0 not sampled. This variable accounts for the temporal misalignment in the CR data by setting *p*_*i,t*_ = 0 when a MAPS location-occasion was not sampled.

We modeled survival and recruitment under the spatio-temporal data misalignment scenario described in section 3.3. We investigated the effect of residual maximum temperature (*TMAX*) on survival and recruitment during the breeding season (May-August). Residual *TMAX* represents local deviations in temperature after removing the broad-scale spatial trend, ensuring that the effect of *TMAX* is not confounded with geographic location and was calculated as the residuals from a linear model of *TMAX* on an intercept and spatial coordinates. Specifically, we considered two cases of residual *TMAX* effect on survival and recruitment: a spatially constant case and a spatially varying case. Spatio-temporal values of *TMAX* were obtained from TerraClimate (Abatzoglou et al., 2018). For each location *i* and year *t, TMAX*_*i,t*_ was defined as the median of monthly mean daily maximum temperatures over the breeding season. For the spatially constant effect, we employed a linear function and for the spatially varying case, we considered an interaction function between spatial coordinates and residual *TMAX*. We centered and standardized all covariates. Section S2.3 in the Supporting Information provides more details on the covariate effect cases, prior settings and MCMC settings.

### 5.3 Results

We applied our sIPM to the BBS and MAPS data considering the two cases of residual *TMAX* effect and models with and without the NNGP on survival and recruitment. In all cases, all parameters of interest converged according to Gelman and Rubin’s convergence diagnostic (Gelman & Rubin, 1992). To select the model that best fitted the data, we performed model selection using posterior predictive loss (*PPL*) (A. Gelfand, 1998). *PPL* is a Bayesian model selection criterion well suited for spatial models. It is a sum of a goodness of fit measure and a penalty that measures how well a model’s predicted data generated from its posterior predictive distribution matches the observed data. Lower PPL indicates better predictive performance. See Supporting Information section S2.5 for mathematical description and Hooten and Hobbs (2015) for more details.

From Table 2, we can see that *PPL* selected the model with spatially varying residual *TMAX* effect and the NNGP as the best one. Thus, there is support of an spatially varying effect of residual *TMAX* on survival and recruitment and for the inclusion of the NNGP to account for residual spatial autocorrelation. In the following we display the results obtained from this model.

**Table 2:** Posterior predictive loss (*PPL*) model selection results for Gray Catbird, comparing models with spatially constant and spatially varying residual maximum temperate (*TMAX*) effects and with and without a nearest-neighbor Gaussian process (NNGP) on survival and recruitment. Spatially constant models assume a single global effect of residual maximum temperature, whereas spatially varying models allow the effect of residual maximum temperature to vary with spatial location. Δ*PPL* = *PPL*_*m*_ − *PPL*^∗^ for model *m* where *PPL*^∗^ is the minimum *PPL* among models considered. Lower *PPL* values indicate better predictive performance.

| Model | Residual $TMAX$ effect | NNGP | $PPL$ | Goodness of fit | Penalty | $\Delta PPL$ |
| --- | --- | --- | --- | --- | --- | --- |
| 1 | Spatially constant | No | 169208.5 | 52330.8 | 116877.6 | 143.9 |
| 2 | Spatially constant | Yes | 169201.9 | 52195.5 | 117006.4 | 137.3 |
| 3 | Spatially varying | No | 169110.9 | 52217.5 | 116893.4 | 46.3 |
| 4 | Spatially varying | Yes | 169064.6 | 51971.7 | 117092.8 | 0 |

Annual estimates of baseline survival, recruitment, relative abundance and growth rate were all temporally heterogeneous (Figure 4), agreeing with the non-spatial analysis results obtained by Ahrestani et al. (2017) and Schaub and Kéry (2022).

**Figure 4.**
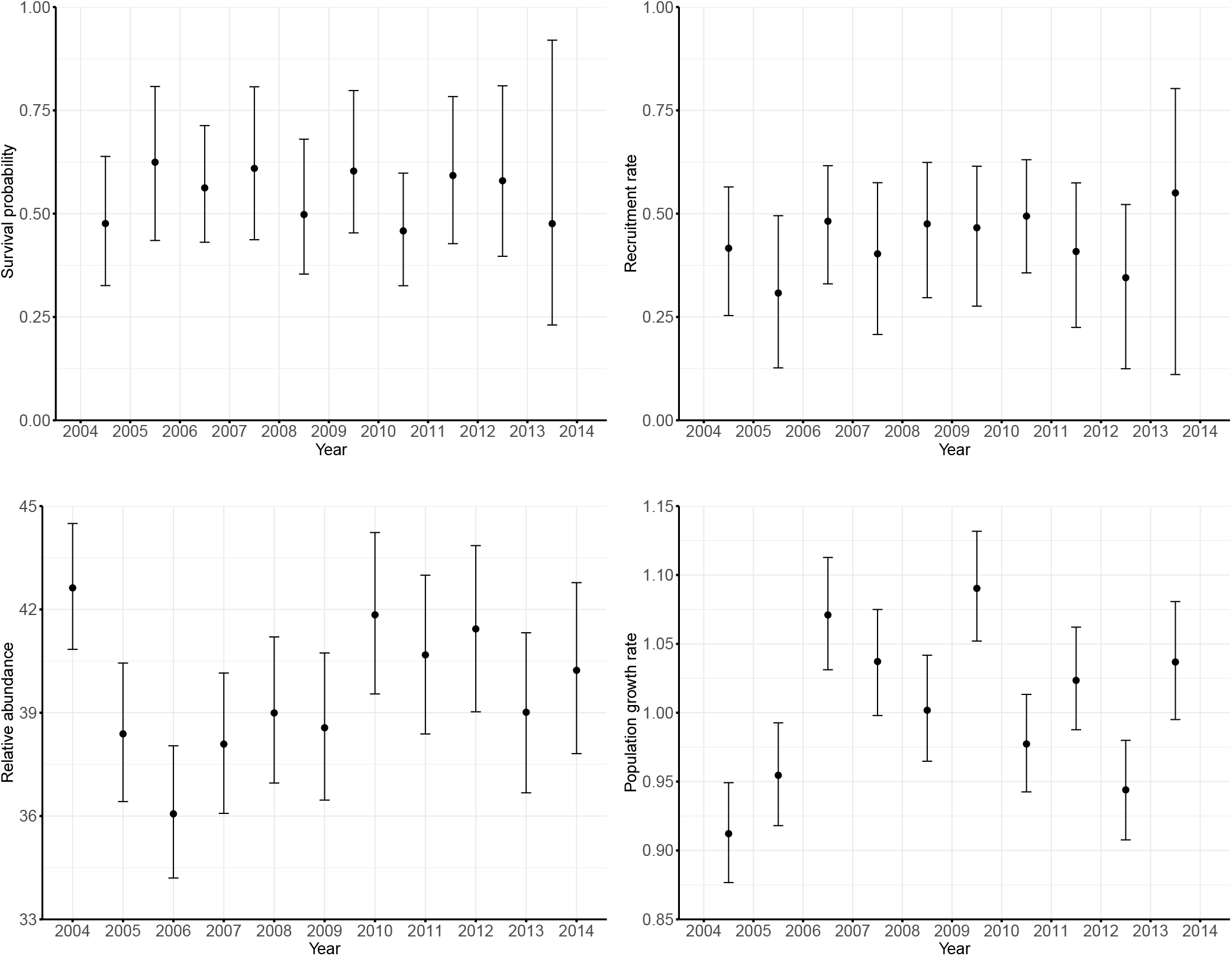
The black dots represent the posterior mean of annual baseline survival, recruitment, relative abundance and population growth rate averaged over space and the vertical bands represent the corresponding 95% posterior credible interval. Survival, recruitment, and population growth rate are defined over the period from year *t* to *t* + 1 and are plotted at the midpoint of each annual interval.

Importantly, Figure 5 highlights the novelty of our modeling framework as it provides continuous spatial predictions of demographic rates and relative abundance across the entire study region, averaged over 2004-2014. Survival and recruitment predictions were made via kriging and predictions of relative abundance and growth rate were produced by Bayesian IDW. In particular, for kriging, we employ algorithm 2 of Finley et al. (2019) to obtain posterior predictive inference on the Nearest Neighbor Gaussian process. For Bayesian IDW, we use the function gstat from the R package `gstat` (Pebesma, 2004) where we set the number of nearest observations to 15. Occasion-specific continuous spatial predictions of these parameters can also be obtained, providing a deeper understanding of how these demographic rates and relative abundance vary across space and time. For simplicity and demonstrative purpose we present continuous spatial predictions across the study region averaged over time.

**Figure 5.**
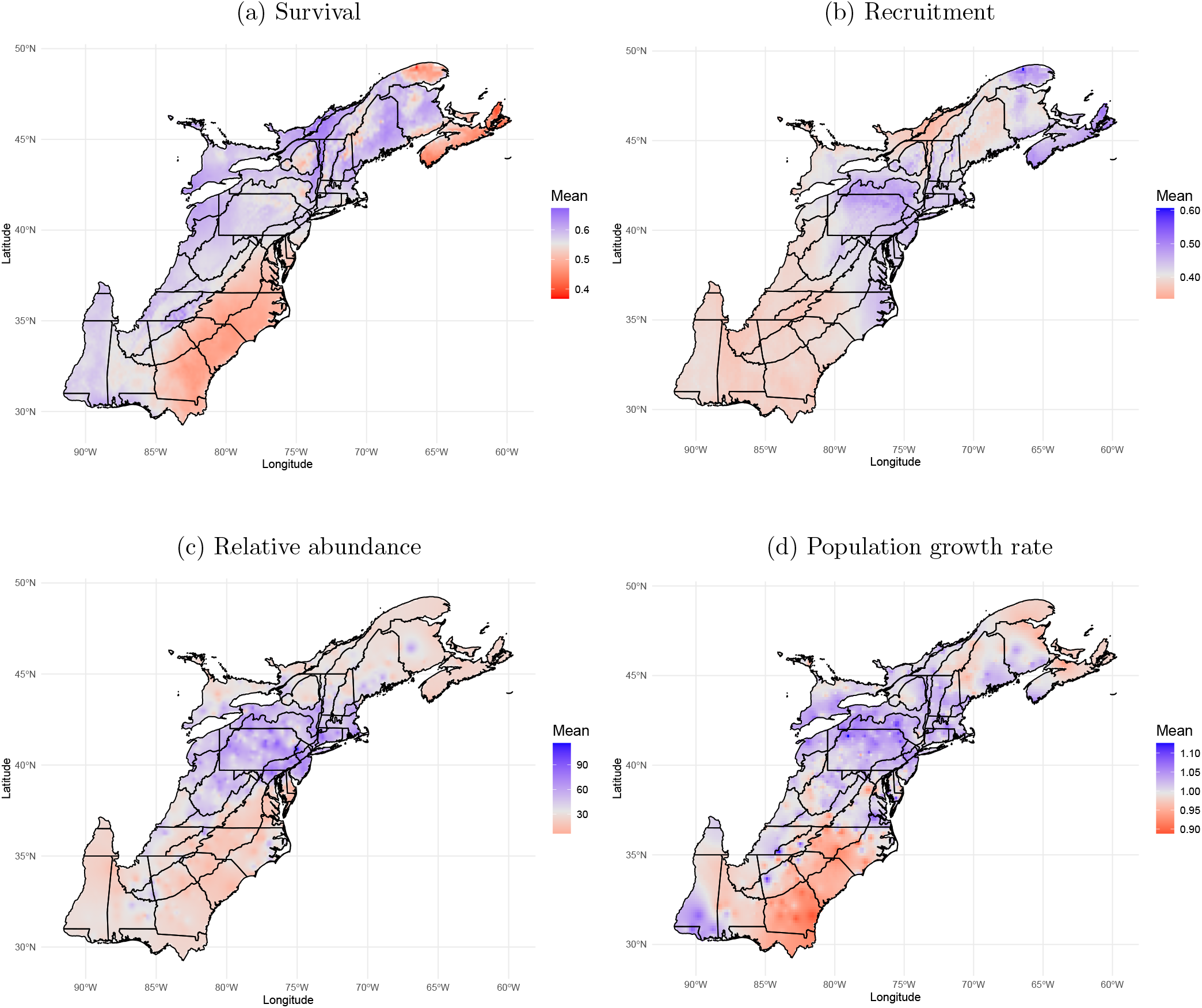
Maps showing the mean predictions of spatial predictions for (a) survival, (b) recruitment, (c) relative abundance and (d) population growth rate of Gray Catbirds across the eastern coast of the United States averaged across 2004-2014.

Spatial predictions revealed pronounced spatial heterogeneity in demographic rates and relative abundance of Gray Catbirds across the eastern coast of the United States during 2004–2014. Survival was highest across the core of the study region, particularly in northeastern areas, and declined toward southern and northern areas. Recruitment exhibited similar broad-scale spatial structure, with higher recruitment concentrated in central areas and consistently lower recruitment across the southern extent of the study region. Relative abundance peaked in central areas where both survival and recruitment were elevated and decreased toward the study region margins. Spatial patterns in growth rate reflected the combined influence of survival and recruitment, with growth rates exceeding one across much of the central and northern areas of the study region and declining below one in southern areas. Across all parameters, spatial prediction uncertainty was modest, lowest in data rich core regions and increased toward the study region boundaries (Figure S6). Moreover, the spatially varying slope of residual *TMAX* revealed complementary patterns between survival and recruitment (Figure 6 and Figure S4). Survival tended to increase with warmer than expected *TMAX* in northern areas but declined under similar conditions in southern areas. In contrast, recruitment exhibited stronger positive effect to warmer than expected *TMAX* in southern areas and weaker effects across northern areas of the study region. Overall, uncertainty in the spatially varying slope was modest, with an increase primarily occurring at the northern and southern edges of the study region (Figure S7). Collectively, these results clearly indicate that survival, recruitment, relative abundance, growth rate and effect of residual *TMAX* were not homogeneous across the study area but instead displayed pronounced spatial heterogeneity.

**Figure 6.**
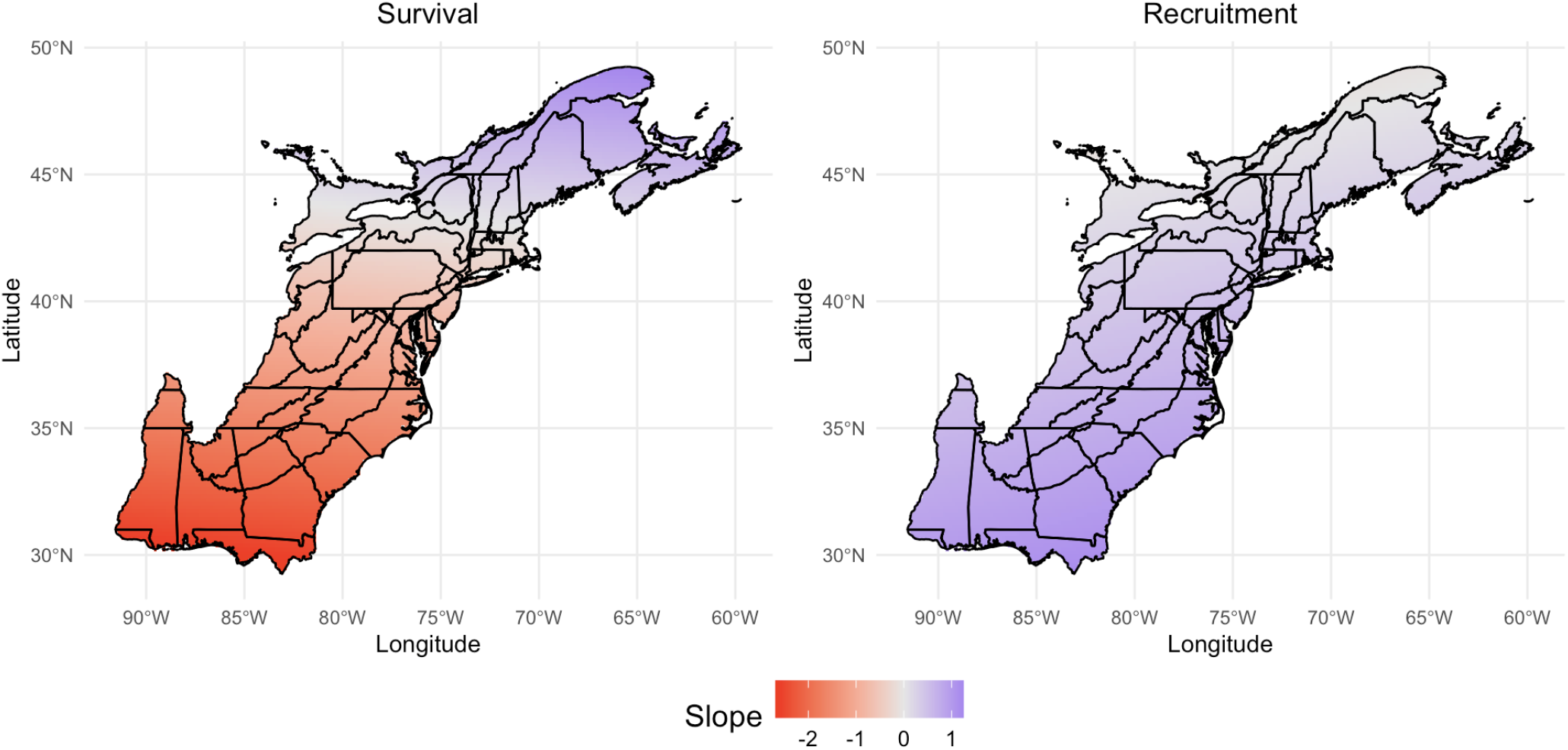
Spatially varying slope of residual maximum temperature for survival and recruitment of Gray Catbirds across the eastern coast of the United States from 2004-2014.

Finally, we evaluated goodness of fit via posterior predictive checks (Hooten & Hobbs, 2015) using the Freeman-Tukey fit statistic. We considered spatial grouping and temporal grouping to better understand how our sIPM fitted the data spatially and temporally, respectively. We further summarized the difference between the actual and predicted Freeman-Tukey fit statistics using Bayesian *p* values, computed separately for the BBS and MAPS datasets. Mathematical details and additional results are provided in the Supporting Information section S2.11. For the BBS count data, Bayesian *p*-values under spatial grouping were approximately 0.5 across all locations, indicating our model adequately captured the spatial variation well. Under temporal grouping, 10 of 11 occasions also suggested good model fit. Similarly, for the MAPS CR data, there was no evidence of lack of fit under temporal grouping, and about 80% of MAPS locations showed no evidence of lack of fit under spatial grouping. Overall, these results indicate that our sIPM adequately fit both data sources across space and time.

## 6 Discussion

Understanding how demographic processes vary across space and time is a central objective to ecology and conservation. With the expansion of spatio-temporal biodiversity monitoring programs, advances in spatial statistics and computation power, IPMs are well suited to provide insights on spatio-temporal variation in demographic processes. However, most IPMs are not spatially explicit and assume that demographic rates are homogeneous across space. We have developed a spatially explicit IPM (sIPM) to estimate and predict spatio-temporal demographic rates, enabling better insights into spatia-temporal population dynamics. Our framework uses a joint likelihood approach to combine population count and CR data. Through the concept of support, it rigorously handles spatial and spatio-temporal data misalignment and accounts for residual spatial autocorrelation efficiently via a nearest-neighbour Gaussian process (NNGP).

We conducted an extensive simulation study to evaluate the performance of our novel modeling framework in the presence of varying degrees of data misalignment, residual spatial autocorrelation and different covariate effects on survival and recruitment. Our results demonstrate that the proposed framework reliably estimates spatio-temporal heterogeneity in demographic rates and abundance across all cases examined. Regression coefficients of survival and recruitment were also accurately recovered whether they were in the spatially constant or spatially varying case, which highlights the framework’s ability to identify key ecological processes. Equally, survival and recruitment demonstrated strong predictive performance in all cases considered. This demonstrates the robustness of our framework under a wide range of conditions.

Using spatio-temporally misaligned population count data from North American Breeding Bird Survey (BBS) and CR data from the Monitoring Avian Productivity and Survivorship program (MAPS), we applied our sIPM to the Gray Catbird across the eastern coast of United States from 2004 to 2014. Results uncovered clear spatio-temporal structuring of demographic rates and relative abundance, with central areas characterized by higher apparent survival, recruitment, relative abundance, and positive population growth rate, and southern areas exhibiting consistently lower demographic performance. Population growth rates emerged from the joint spatial structure of survival and recruitment, indicating that demographic processes covary spatially rather than operating uniformly across the study region. This spatial concordance is consistent with the potential emergence of source–sink dynamics (Pulliam, 1988), where central areas may serve as as demographic sources and southern peripheral areas as demographic sinks. These insights are critical for spatially informing targeted conservation and management strategies. Parameter uncertainty was modest, further supporting the robustness of this framework. Together, these results illustrate the value of our sIPM to uncover important spatio-temporal heterogeneity in demographic rates and relative abundance that would be obscured under traditional, non-spatial IPMs.

Notably, our analysis also provided new insights into how residual maximum temperature (*TMAX*) influenced Gray Catbird survival and recruitment during the breeding season, with effects varying geographically across the study region. By incorporating an interaction between residual *TMAX* and spatial coordinates, we found complementary spatial patterns in survival and recruitment: in northern areas, warmer than expected *TMAX* was associated with increased survival but weaker recruitment, whereas southern regions exhibited the opposite pattern. Together, these results highlight spatial heterogeneity not only in demographic rates but also in climate sensitivity and underscores the need for models that allow environmental drivers to act in spatially heterogeneous ways. However, this interaction function assumes a linear spatial trend in residual *TMAX* with respect to spatial coordinates, which may oversimplify reality and potentially lead to misleading inference. To address this, future work can incorporate spatial varying coefficients (SVCs; Doser et al. (2025) and Finley (2011)), a flexible approach for modeling non-linear spatially varying covariate effects in our sIPM. One promising avenue is to leverage the nearest-neighbor Gaussian process(NNGP) to model the spatially varying effect of a covariate, making the integration of SVCs feasible, as demonstrated in R packages such as spOccupancy (Doser et al., 2022) and spAbundance (Doser et al., 2024). It is worth noting that our framework can estimate absolute abundance once the sampling area is known. However, predictions via IDW of absolute abundance cannot be made at a finer resolution than that estimated at as absolute abundance is inherently dependent on the spatial resolution. This importantly, does not affect the prediction of survival, recruitment and population growth rate as by definition these are independent of the spatial resolution.

Our sIPM can be readily applied to other species whose life-cycle can be represented by a single age class. For species with more complex life-cycles that include age-dependent survival or delayed start of reproduction, the population model needs to be changed. Consequently, extending our sIPM to include age-dependence is an important next step. Another important extension would be the explicit incorporation of dispersal. Our framework produces crucial estimates of apparent survival and recruitment, but these are confounded with dispersal. By explicitly accounting for dispersal would enable the estimation of true survival and the decomposition of recruitment into local recruitment and immigration, leading to more robust and ecologically informative inference. These extensions would require integrating complementary data sources such as productivity data, age-dependent CR data, or tracking data into our sIPM.

In the NNGP, we use an exponential covariance function to determine the similarity between locations. Although flexible and suitable to a wide range of cases there are caveats to this covariance function. Namely, it assumes stationary, i.e, covariance depends only on the Euclidean distance between locations.

This assumption may be violated when domain knowledge suggests that the underlying process varies more rapidly in some regions of the spatial domain than in others, independent of Euclidean distance. For instance, environmental variables are often much less smooth in mountainous areas than in flat regions. In such cases, non-stationary GP’s (Paciorek & Schervish, 2003) should be used as they adapt to changes in the smoothness of the function of interest. In addition, we assume a separable GP, meaning space and time are treated as independent. Consequently, residual spatial autocorrelation is assumed constant across time. Non-separable GPs (Datta, Banerjee, Finley, Hamm, & Schaap, 2016) can be used to relax this assumption and to capture space-time interactions. Separable GPs offer simplicity and computational efficiency, whereas non-separable GPs offer greater flexibility for modeling interactions between space and time, but are generally more computationally intensive. Therefore, future work can be focused on extending our sIPM to include non-stationary and non-separable GPs.

With all the possible avenues of model extension, sIPMs complexity grows and computational efficiency emerges as a key issue. Future work will therefore need to explore computational solutions to make these extensions feasible for real-world applications. One promising direction involves optimizing the MCMC sampling of NNGP spatial terms, using techniques like the surrogate data slice sampling (Murray & Adams, 2010) modified by Wright and Hooten (2025), which can significantly reduce the computational cost of high-dimensional NNGPs. Furthermore, transitioning toward likelihood-free inference, including approximate Bayesian computation (ABC) (Sisson et al., 2018) or Neural Bayes Estimators (Sainsbury-Dale et al., 2024), offers an appealing path forward. These approaches have demonstrated the ability to provide rapid, accurate parameter estimation for complex models, potentially bypassing the intensive run-times currently required for spatially explicit IPMs.

In conclusion, by estimating spatio-temporal demographic rates at their native spatial supports and predicting these rates at high spatial resolutions, our sIPM provides the critical insights needed to address a long standing challenge in ecology: understanding spatio-temporal variation in demographic processes. This enables the move beyond one-size-fits-all conservation and the development of strategies that are not only effective, but are specifically optimized to the unique needs of a population in specific locations and times.

## Author Contributions

Fabian R. Ketwaroo and Michael Schaub conceived the idea. Fabian R. Ketwaroo designed and implemented the model, analysed the data and led writing. Michael Schaub and Matia. H. Muller contributed to writing. James F. Saracoo reviewed the results and the manuscript. All authors contributed critically to the drafts and gave final approval for publication.

## Statement on inclusion

We have considered equity, diversity and inclusion in the development and presentation of this work. We aimed to cite a diverse and representative body of literature and to develop a modeling framework that is broadly accessible and reproducible. The full model specification and implementation details are provided to facilitate use by researchers across ecological contexts.

## Acknowledgements

Financial support was provided by grant no. 215689 (to Michael Schaub) from the Swiss National Science Foundation. Thanks to Danielle Kaschube for assisting with Monitoring Avian Productivity and Survivorship program (MAPS) data preparation.

## Data Availability

All data sources used in this study are properly cited in the manuscript. Gray Catbird observations from the BBS (North American Breeding Bird Survey) are publicly available and can be accessed via the R package `bbsBayes2` (Edwards et al., 2023). Raw MAPS (Monitoring Avian Productivity and Survivorship program) data used to construct capture histories are publicly available through the MAPS Data Exploration Tool (Institute for Bird Populations, 2023). R code and data supporting manuscript available at https://doi.org/10.5281/zenodo.18846082

## Supporting Information

Supporting Information includes the following sections: S1. Simulation study and S2. Case study: Gray Catbird.

## References

Abadi, F., Gimenez, O., Ullrich, B., Arlettaz, R., & Schaub, M. (2010). Estimation of immigration rate using integrated population models. Journal of Applied Ecology, 47 (2), 393–400. 10.1111/j.1365-2664.2010.01789.x

Abatzoglou, J. T., Dobrowski, S. Z., Parks, S. A., & Hegewisch, K. C. (2018). Terraclimate, a high-resolution global dataset of monthly climate and climatic water balance from 1958–2015. Scientific data, 5 (1), 1–12.

Ahrestani, F. S., Saracco, J. F., Sauer, J. R., Pardieck, K. L., & Royle, J. A. (2017). An integrated population model for bird monitoring in North America. Ecological Applications, 27 (3), 916–924. 10.1002/eap.1493

Banerjee, S., & Fuentes, M. (2012). Bayesian modeling for large spatial datasets. Wiley Interdisciplinary Reviews: Computational Statistics, 4 (1), 59–66.

Besbeas, P., Freeman, S. N., Morgan, B. J., & Catchpole, E. (2002). Integrating mark–recapture–recovery and census data to estimate animal abundance and demographic parameters. Biometrics, 58 (3), 540–547.

Bosley, K. M., Schueller, A. M., Goethel, D. R., Hanselman, D. H., Fenske, K. H., Berger, A. M., Deroba, J. J., & Langseth, B. J. (2022). Finding the perfect mismatch: Evaluating misspecification of population structure within spatially explicit integrated population models. Fish and Fisheries, 23 (2), 294–315.

Burrough, P. A., McDonnell, R. A., & Lloyd, C. D. (2015). Principles of geographical information systems. Oxford university press.

Cave, V. M., King, R., & Freeman, S. N. (2010). An integrated population model from constant effort bird-ringing data. Journal of Agricultural, Biological, and Environmental Statistics, 15 (1), 119–137.

Cole, D. J., & McCrea, R. S. (2016). Parameter redundancy in discrete state-space and integrated models. Biometrical Journal, 58 (5), 1071–1090.

Coulson, T., Catchpole, E. A., Albon, S. D., Morgan, B. J., Pemberton, J., Clutton-Brock, T. H., Crawley, M., & Grenfell, B. (2001). Age, sex, density, winter weather, and population crashes in soay sheep. Science, 292 (5521), 1528–1531.

Cressie, N. (1990). The origins of kriging. Mathematical geology, 22 (3), 239–252.

Cressie, N. (2015). Statistics for spatial data. John Wiley & Sons.

Datta, A., Banerjee, S., Finley, A. O., & Gelfand, A. E. (2016). Hierarchical nearest-neighbor Gaussian process models for large geostatistical datasets. Journal of the American Statistical Association, 111 (514), 800–812.

Datta, A., Banerjee, S., Finley, A. O., Hamm, N. A., & Schaap, M. (2016). Nonseparable dynamic nearest neighbor Gaussian process models for large spatio-temporal data with an application to particulate matter analysis. The annals of applied statistics, 10 (3), 1286.

David Jr Ziolkowski, U., Lutmerding, M., English, W., Hudson, M.-A., et al. (2025). North American Breeding Bird Survey Dataset 1966-2023.

Desante, D. F., Burton, K. M., Saracco, J. F., & Walker, B. L. (1995). Productivity indices and survival rate estimates from MAPS, a continent-wide programme of constant-effort mist-netting in North America. Journal of Applied Statistics, 22 (5-6), 935–948.

DeSante, D. F., & Kaschube, D. R. (2009). The monitoring avian productivity and survivorship (maps) program 2004, 2005, and 2006 report. Bird Populations, 9, 86–169.

DeSante, D. F., Saracco, J. F., O’grady, D. R., Burton, K. M., & Walker, B. L. (2004). Methodological considerations of the monitoring avian productivity and survivorship (maps) program. Studies in Avian Biology, 28–45.

de Valpine, P., Turek, D., Paciorek, C. J., Anderson-Bergman, C., Lang, D. T., & Bodik, R. (2017). Programming with models: Writing statistical algorithms for general model structures with NIMBLE. Journal of Computational and Graphical Statistics, 26 (2), 403–413.

Doser, J. W., Finley, A. O., Kéry, M., & Zipkin, E. F. (2022). spOccupancy: An R package for single-species, multi-species, and integrated spatial occupancy models. Methods in Ecology and Evolution, 13 (8), 1670–1678.

Doser, J. W., Finley, A. O., Kéry, M., & Zipkin, E. F. (2024). spAbundance: An R package for single-species and multi-species spatially explicit abundance models. Methods in Ecology and Evolution, 15 (6), 1024–1033.

Doser, J. W., Finley, A. O., Saunders, S. P., Kéry, M., Weed, A. S., & Zipkin, E. F. (2025). Modeling complex species-environment relationships through spatially-varying coefficient occupancy models. Journal of Agricultural, Biological and Environmental Statistics, 30 (1), 146–171.

Dungan, J. L., Perry, J., Dale, M., Legendre, P., Citron-Pousty, S., Fortin, M.-J., Jakomulska, A., Miriti, M., & Rosenberg, M. (2002). A balanced view of scale in spatial statistical analysis. Ecography, 25 (5), 626–640.

Edwards, B., Smith, A., & LaZerte, S. (2023). bbsBayes2: Hierarchical Bayesian analysis of North American BBS data. R package.

Finley, A. O. (2011). Comparing spatially-varying coefficients models for analysis of ecological data with non-stationary and anisotropic residual dependence. Methods in ecology and evolution, 2 (2), 143–154.

Finley, A. O., Datta, A., Cook, B. D., Morton, D. C., Andersen, H. E., & Banerjee, S. (2019). Efficient algorithms for Bayesian Nearest neighbor Gaussian Processes. Journal of Computational and Graphical Statistics, 28 (2), 401–414. 10.1080/10618600.2018.1537924

Fournier, D., & Archibald, C. P. (1982). A general theory for analyzing catch at age data. Canadian Journal of Fisheries and Aquatic Sciences, 39 (8), 1195–1207.

Frost, F., McCrea, R., King, R., Gimenez, O., & Zipkin, E. (2023). Integrated population models: Achieving their potential. Journal of Statistical Theory and Practice, 17 (1), 6. 10.1007/s42519-022-00302-7

Gaillard, J.-M., Festa-Bianchet, M., Yoccoz, N. G., Loison, A., & Toıgo, C. (2000). Temporal variation in fitness components and population dynamics of large herbivores. Annual Review of ecology and Systematics, 31 (1), 367–393.

Gelfand, A. (1998). Model choice: A minimum posterior predictive loss approach. Biometrika, 85 (1), 1–11. 10.1093/biomet/85.1.1

Gelfand, A. E., Zhu, L., & Carlin, B. P. (2001). On the change of support problem for spatio-temporal data. Biostatistics, 2 (1), 31–45.

Gelman, A., & Rubin, D. B. (1992). Inference from iterative simulation using multiple sequences. Statistical science, 7 (4), 457–472.

Gotway, C. A., & Young, L. J. (2002). Combining incompatible spatial data. Journal of the American Statistical Association, 97 (458), 632–648.

Haining, R. P., & Li, G. (2020). Modelling spatial and spatial-temporal data: A Bayesian approach. CRC Press.

Holt, R. D. (1984). Spatial heterogeneity, indirect interactions, and the coexistence of prey species. The American Naturalist, 124 (3), 377–406.

Hooten, M. B., & Hobbs, N. T. (2015). A guide to Bayesian model selection for ecologists. Ecological Monographs, 85 (1), 3–28. 10.1890/14-0661.1

Hostetler, J. A., & Chandler, R. B. (2015). Improved state-space models for inference about spatial and temporal variation in abundance from count data. Ecology, 96 (6), 1713–1723.

Institute for Bird Populations. (2023). MAPS Data Exploration Tool: Downloaded MAPS data [24.02.2026]. https://ibp-maps-data-exploration-tool.org

Lebreton, J.-D., Burnham, K. P., Clobert, J., & Anderson, D. R. (1992). Modeling survival and testing biological hypotheses using marked animals: A unified approach with case studies. Ecological monographs, 62 (1), 67–118.

Link, W. A., & Sauer, J. R. (2002). A hierarchical analysis of population change with application to cerulean warblers. Ecology, 83 (10), 2832–2840.

Michener, W. K., & Jones, M. B. (2012). Ecoinformatics: Supporting ecology as a data-intensive science. Trends in ecology & evolution, 27 (2), 85–93.

Murray, I., & Adams, R. P. (2010). Slice sampling covariance hyperparameters of latent Gaussian models. Advances in neural information processing systems, 23.

Pacifici, K., Reich, B. J., Miller, D. A. W., & Pease, B. S. (2019). Resolving misaligned spatial data with integrated species distribution models. Ecology, 100 (6), e02709. 10.1002/ecy.2709

Paciorek, C., & Schervish, M. (2003). Nonstationary covariance functions for Gaussian process regression. Advances in neural information processing systems, 16.

Pavliotis, G. A. (2014). Stochastic processes and applications. Texts in applied mathematics, 60.

Pebesma, E. J. (2004). Multivariable geostatistics in S: The gstat package. Computers Geosciences, 30, 683–691. 10.1016/j.cageo.2004.03.012

Pradel, R., Hines, J. E., Lebreton, J.-D., & Nichols, J. D. (1997). Capture-recapture survival models taking account of transients. Biometrics, 60–72.

Prochazka, B. G., Coates, P. S., O’Neil, S. T., Espinosa, S. P., & Aldridge, C. L. (2024). Geographic principles applied to population dynamics: A spatially interpolated integrated population model. Methods in Ecology and Evolution, 15 (8), 1394–1407. 10.1111/2041-210X.14334

Pulliam, H. R. (1988). Sources, sinks, and population regulation. The American Naturalist, 132 (5), 652–661.

Sainsbury-Dale, M., Zammit-Mangion, A., & Huser, R. (2024). Likelihood-free parameter estimation with neural Bayes estimators. The American Statistician, 78 (1), 1–14.

Sauer, J., Hines, J., Fallon, J., Pardieck, K., Ziolkowski Jr, D., & Link, W. (2014). The North American breeding bird survey, results and analysis 1966–2013. Version 01.30. 2015. US Geological Survey Patuxent Wildlife Research Center, Laurel, Maryland, USA.

Saunders, S. P., Farr, M. T., Wright, A. D., Bahlai, C. A., Ribeiro Jr, J. W., Rossman, S., Sussman, A. L., Arnold, T. W., & Zipkin, E. F. (2019). Disentangling data discrepancies with integrated population models. Ecology, 100 (6), e02714.

Schaub, M., & Abadi, F. (2011). Integrated population models: A novel analysis framework for deeper insights into population dynamics. Journal of Ornithology, 152 (Suppl 1), 227–237.

Schaub, M., & Kéry, M. (2022). Integrated population models: Theory and ecological applications with R and JAGS. Academic Press.

Sisson, S. A., Fan, Y., & Beaumont, M. (2018). Handbook of approximate Bayesian computation. CRC press.

Tobler, W. R. (1970). A computer movie simulating urban growth in the detroit region. Economic geography, 46 (up1), 234–240.

Vecchia, A. V. (1988). Estimation and model identification for continuous spatial processes. Journal of the Royal Statistical Society Series B: Statistical Methodology, 50 (2), 297–312.

Williams, C. K., & Rasmussen, C. E. (2006). Gaussian processes for machine learning (Vol. 2). MIT press Cambridge, MA.

Wright, W. J., & Hooten, M. B. (2025). Continuous-space occupancy models. Biometrics, 81 (2), ujaf055.

Zhao, Q. (2020). On the sampling design of spatially explicit integrated population models.

Zipkin, E. F., Zylstra, E. R., Wright, A. D., Saunders, S. P., Finley, A. O., Dietze, M. C., Itter, M. S., & Tingley, M. W. (2021). Addressing data integration challenges to link ecological processes across scales. Frontiers in Ecology and the Environment, 19 (1), 30–38. 10.1002/fee.2290

