## Supporting information for "An Integrated Population Model to Incorporate Spatio-Temporal Heterogeneity in Demographic Rates"

#### S1 Simulation Study

To evaluate the simulation studies, we use the mean of the posterior means, relative bias $\big(\text{RB} = (\bar{v} - v)/v\big)$, root mean square error $\big(\text{RMSE} = \sqrt{\frac{1}{n}\sum_{i=1}^{n}(\bar{v}_i - v)^2}\big)$ where $n$ is the simulation runs, $v$ is the true parameter value and $\bar{v}_i$ is the $i^{th}$ estimated posterior mean. We also use the 95% posterior credible interval coverage (Cov) to summarise parameter estimation across simulation runs. For predictions, we use mean absolute percentage error $(\text{MAPE} = \frac{\sum_{i=1}^{m}|\frac{v-\tilde{v}}{v}|\times 100}{m})$ where $\tilde{v}$ is the predicted value. For both estimation and prediction use also compute Monte Carlo standard error of RB $(\text{MCSE(RB)} = \sqrt{Var(\text{RB})})$.

##### S1.1 Settings

Let $\boldsymbol{X}_t$ be a spatially correlated covariate at sampling occasion $t$ such that $\boldsymbol{X}_t \sim GP(0, \boldsymbol{K})$ where $\boldsymbol{K}$ is an exponential covariance function with length scale = 0.4 and variance = 0.2. Table S1 shows the parameter values of occasion-specific baseline recapture probability $(\alpha_t^p)$, spatially constant regression effects $(\beta^f, \beta^\phi)$ of survival and recruitment, spatially varying regression effects $(\beta_{1:5}^f, \beta_{1:5}^\phi)$ of survival and recruitment, NNGP parameters $(\sigma_f^2, \rho^f, \sigma_\phi^2, \rho^\phi)$ of survival and recruitment, occasion-specific baseline survival $(\alpha_t^\phi)$ and occasion-specific baseline recruitment $(\alpha_t^f)$ used to simulate data. Table S2 gives the prior settings used in the simulation study.

For the cases with a spatially constant covariate effect on survival and recruitment: 2 MCMC chains of 85000 iterations, 60000 burn-in were performed each run. In these cases, parameters in Table S1 were thinned by 10 while survival, recruitment, population abundance and population growth rate where thinned by 20. For the cases with spatially varying covariate effect on survival and recruitment: 2 MCMC chains of 125000 iterations, 80000 burn-in were performed each run. Likewise, parameters in Table S1 were thinned by 15 while survival, recruitment, population abundance and population growth rate where thinned by 25. In all cases, all parameters converged according to Gelman and Rubin's convergence diagnostic (Gelman & Rubin, 1992).

| Covariate effect | Parameter | Value |
|---|---|---|
| Spatially constant | CR releases | Discrete Uniform(20, 50) |
| | Initial Abundance | Discrete Uniform(80, 120) |
| | $\alpha_t^p$ | logit(Uniform(0.4, 0.7)) |
| | $\beta^f$ | $-0.2$ |
| | $\beta^\phi$ | $-0.15$ |
| | $\rho_f$ | 0.35 |
| | $\sigma_f^2$ | 0.1 |
| | $\rho_\phi$ | 0.3 |
| | $\sigma_\phi^2$ | 0.2 |
| | $\alpha_t^\phi$ | logit(Uniform(0.5, 0.85)) |
| | $\alpha_t^f$ | log(1- $\text{logit}^{-1}(\alpha_t^\phi)$ ) |
| Spatially varying | CR releases | Discrete Uniform(20, 50) |
| | Initial Abundance | Discrete Uniform(80, 120) |
| | $\alpha_t^p$ | logit(Uniform(0.4, 0.7)) |
| | $\beta_1^f$ | $-0.2$ |
| | $\beta_2^f$ | 0.1 |
| | $\beta_3^f$ | $-0.4$ |
| | $\beta_4^f$ | 0.2 |
| | $\beta_5^f$ | 0.2 |
| | $\beta_1^\phi$ | $-0.15$ |
| | $\beta_2^\phi$ | 0.1 |
| | $\beta_3^\phi$ | 0.5 |
| | $\beta_4^\phi$ | $-0.2$ |
| | $\beta_5^\phi$ | $-0.5$ |
| | $\rho_f$ | 0.35 |
| | $\sigma_f^2$ | 0.1 |
| | $\rho_\phi$ | 0.3 |
| | $\sigma_\phi^2$ | 0.2 |
| | $\alpha_t^\phi$ | logit(Uniform(0.5, 0.85)) |
| | $\alpha_t^f$ | log(1- $\text{logit}^{-1}(\alpha_t^\phi)$ ) |

Table S1: Data generating values of occasion-specific baseline recapture probability ($\alpha_t^p$), spatially constant regression effects ($\beta^f, \beta^\phi$) of survival and recruitment, spatially varying regression effects ($\beta_{1:5}^f, \beta_{1:5}^\phi$) of survival and recruitment, Nearest Neighbor Gaussian Process parameters ($\sigma_f^2$, $\rho^f$, $\sigma_\phi^2, \rho^\phi$) of survival and recruitment, occasion-specific baseline survival ($\alpha_t^\phi$) and occasion-specific baseline recruitment ($\alpha_t^f$).

| Parameter | Prior |
|---|---|
| Initial abundance | Discrete Uniform (50, 150) |
| $\alpha_t^p$ | Normal (0, $\sigma^2 = 2$) |
| $\beta^f$ | Normal(0, $\sigma^2$= 2) |
| $\beta^\phi$ | Normal(0, $\sigma^2$= 2) |
| $\beta_1^f$ | Normal(0, $\sigma^2$= 2) |
| $\beta_2^f$ | Normal(0, $\sigma^2$= 2) |
| $\beta_3^f$ | Normal(0, $\sigma^2$= 2) |
| $\beta_4^f$ | Normal(0, $\sigma^2$= 2) |
| $\beta_5^f$ | Normal(0, $\sigma^2$= 2) |
| $\beta_1^\phi$ | Normal(0, $\sigma^2$= 2) |
| $\beta_2^\phi$ | Normal(0, $\sigma^2$= 2) |
| $\beta_3^\phi$ | Normal(0, $\sigma^2$= 2) |
| $\beta_4^\phi$ | Normal(0, $\sigma^2$= 2) |
| $\beta_5^\phi$ | Normal(0, $\sigma^2$= 2) |
| $\rho_f$ | Uniform($\frac{dmin}{3}$, $\frac{dmax}{3}$) |
| $\sigma_f^2$ | Inverse Gamma(3,1) |
| $\rho_\phi$ | Uniform($\frac{dmin}{3}$, $\frac{dmax}{3}$) |
| $\sigma_\phi^2$ | Inverse Gamma(3,1) |
| $\alpha_t^\phi$ | Normal (0, $\sigma^2 = 2$) |
| $\alpha_t^f$ | Normal (0, $\sigma^2 = 2$) |

Table S2: Prior settings of occasion-specific baseline recapture probability ($\alpha_t^p$), spatially constant regression effects ($\beta^f, \beta^\phi$) of survival and recruitment, spatially varying regression effects ($\beta_{1:5}^f, \beta_{1:5}^\phi$) of survival and recruitment, Nearest Neighbor Gaussian Process parameters ($\sigma_f^2$, $\rho^f$, $\sigma_\phi^2, \rho^\phi$) of survival and recruitment, occasion-specific baseline survival ($\alpha_t^\phi$) and occasion-specific baseline recruitment ($\alpha_t^f$). $dmin, dmax$ denote minimum and maximum Euclidean distance.

#### S1.2 Results

| Covariate effect | Case | Parameter | Mean RB | 2.5% RB | 97.5% RB | MCSE(RB) | RMSE | Cov |
|---|---|---|---|---|---|---|---|---|
| Spatially constant | PA | Survival | 0.003 | -0.076 | 0.095 | 0.007 | 0.025 | 0.950 |
| | | Recruitment | 0.003 | -0.175 | 0.221 | 0.016 | 0.034 | 0.957 |
| | | Abundance | 0.005 | -0.132 | 0.155 | 0.002 | 7.084 | 0.970 |
| | | Growth rate | 0.005 | -0.125 | 0.155 | 0.001 | 0.067 | 0.978 |
| Spatially constant | SM | Survival | 0.008 | -0.117 | 0.163 | 0.015 | 0.041 | 0.943 |
| | | Recruitment | 0.002 | -0.228 | 0.296 | 0.032 | 0.046 | 0.951 |
| | | Abundance | 0.005 | -0.132 | 0.156 | 0.002 | 7.102 | 0.970 |
| | | Growth rate | 0.005 | -0.125 | 0.155 | 0.001 | 0.068 | 0.979 |
| Spatially constant | STM:Low | Survival | 0.010 | -0.119 | 0.170 | 0.015 | 0.042 | 0.943 |
| | | Recruitment | -0.000 | -0.241 | 0.309 | 0.032 | 0.048 | 0.950 |
| | | Abundance | 0.006 | -0.144 | 0.174 | 0.002 | 7.889 | 0.970 |
| | | Growth rate | 0.005 | -0.129 | 0.164 | 0.002 | 0.070 | 0.980 |
| Spatially constant | STM:High | Survival | 0.010 | -0.128 | 0.188 | 0.019 | 0.047 | 0.942 |
| | | Recruitment | -0.001 | -0.272 | 0.353 | 0.042 | 0.055 | 0.944 |
| | | Abundance | 0.011 | -0.181 | 0.244 | 0.004 | 10.714 | 0.972 |
| | | Growth rate | 0.006 | -0.141 | 0.185 | 0.003 | 0.078 | 0.984 |
| Spatially varying | STM:Low | Survival | 0.009 | -0.144 | 0.207 | 0.013 | 0.040 | 0.944 |
| | | Recruitment | 0.004 | -0.230 | 0.299 | 0.027 | 0.047 | 0.953 |
| | | Abundance | 0.007 | -0.146 | 0.181 | 0.002 | 8.507 | 0.971 |
| | | Growth rate | 0.006 | -0.133 | 0.170 | 0.002 | 0.071 | 0.979 |
| Spatially varying | STM:High | Survival | 0.014 | -0.148 | 0.228 | 0.013 | 0.043 | 0.948 |
| | | Recruitment | -0.011 | -0.272 | 0.313 | 0.032 | 0.054 | 0.946 |
| | | Abundance | 0.012 | -0.185 | 0.251 | 0.004 | 12.516 | 0.972 |
| | | Growth rate | 0.006 | -0.145 | 0.195 | 0.002 | 0.079 | 0.981 |

Table S3: Mean of the mean relative bias (RB), mean 2.5% RB, mean 97.5% RB, Monte Carlo standard error of mean RB (MCSE(RB)), mean root mean square error (RMSE) and mean 95% posterior credible interval coverage (Cov) across locations and occasion of survival, recruitment, abundance and growth rate for data integration cases: perfect data alignment (PA), spatial data misalignment (SM), spatial data misalignment with low temporal data misalignment (STM:Low) and spatial data misalignment with high temporal data misalignment (STM:High) with spatially constant and spatially varying covariate effects on survival and recruitment.

| Covariate effect | Case | Parameter | RB | MCSE(RB) | RMSE | Cov |
|---|---|---|---|---|---|---|
| Spatially constant | PA | $\alpha_1^p$ | -0.107 | 0.471 | 0.019 | 0.950 |
| | | $\alpha_2^p$ | 0.009 | 0.179 | 0.015 | 0.970 |
| | | $\alpha_3^p$ | 0.057 | 0.568 | 0.019 | 0.930 |
| | | $\alpha_4^p$ | -0.018 | 0.207 | 0.019 | 0.940 |
| | | $\alpha_5^p$ | 0.193 | 2.221 | 0.052 | 0.950 |
| | | $\sigma_f^2$ | 0.047 | 0.129 | 0.011 | 1.000 |
| | | $\sigma_\phi^2$ | 0.074 | 0.113 | 0.022 | 1.000 |
| | | $\beta^f$ | -0.028 | 0.147 | 0.024 | 0.980 |
| | | $\beta^\phi$ | 0.048 | 0.221 | 0.025 | 0.940 |
| | | $\rho^f$ | 0.139 | 0.029 | 0.048 | 1.000 |
| | | $\rho^\phi$ | 0.126 | 0.103 | 0.043 | 0.980 |
| | | $\alpha_1^f$ | 0.025 | 0.029 | 0.037 | 0.870 |
| | | $\alpha_2^f$ | -0.002 | 0.028 | 0.026 | 0.960 |
| | | $\alpha_3^f$ | 0.002 | 0.028 | 0.024 | 0.940 |
| | | $\alpha_4^f$ | -0.004 | 0.026 | 0.025 | 0.950 |
| | | $\alpha_5^f$ | 0.012 | 0.049 | 0.047 | 0.980 |
| | | $\alpha_1^\phi$ | 0.009 | 0.165 | 0.024 | 0.930 |
| | | $\alpha_2^\phi$ | -0.023 | 0.352 | 0.023 | 0.960 |
| | | $\alpha_3^\phi$ | 0.167 | 1.385 | 0.022 | 0.970 |
| | | $\alpha_4^\phi$ | -0.085 | 1.049 | 0.029 | 0.980 |
| | | $\alpha_5^\phi$ | -0.077 | 0.853 | 0.077 | 0.970 |
| Spatially constant | SM | $\alpha_1^p$ | -0.075 | 0.756 | 0.045 | 0.940 |
| | | $\alpha_2^p$ | 0.061 | 0.401 | 0.038 | 0.950 |
| | | $\alpha_3^p$ | 0.199 | 1.481 | 0.038 | 0.940 |
| | | $\alpha_4^p$ | -0.008 | 0.451 | 0.044 | 0.940 |
| | | $\alpha_5^p$ | 0.121 | 1.165 | 0.087 | 0.940 |
| | | $\sigma_f^2$ | 0.116 | 0.139 | 0.015 | 1.000 |
| | | $\sigma_\phi^2$ | 0.035 | 0.162 | 0.025 | 0.970 |
| | | $\beta^f$ | -0.035 | 0.257 | 0.041 | 0.950 |
| | | $\beta^\phi$ | 0.027 | 0.467 | 0.056 | 0.950 |
| | | $\rho^f$ | 0.121 | 0.032 | 0.042 | 1.000 |
| | | $\rho^\phi$ | 0.173 | 0.083 | 0.053 | 0.990 |
| | | $\alpha_1^f$ | 0.032 | 0.043 | 0.048 | 0.890 |
| | | $\alpha_2^f$ | 0.005 | 0.046 | 0.043 | 0.970 |
| | | $\alpha_3^f$ | 0.008 | 0.048 | 0.042 | 0.930 |
| | | $\alpha_4^f$ | 0.005 | 0.042 | 0.042 | 0.980 |
| | | $\alpha_5^f$ | 0.021 | 0.079 | 0.074 | 0.960 |
| | | $\alpha_1^\phi$ | 0.055 | 0.426 | 0.048 | 0.950 |
| | | $\alpha_2^\phi$ | -0.109 | 0.707 | 0.054 | 0.950 |
| | | $\alpha_3^\phi$ | 0.039 | 0.389 | 0.052 | 0.920 |
| | | $\alpha_4^\phi$ | 0.013 | 0.227 | 0.066 | 0.950 |
| | | $\alpha_5^\phi$ | -0.141 | 2.266 | 0.124 | 0.950 |
| Spatially constant | STM:Low | $\alpha_1^p$ | -0.055 | 0.844 | 0.052 | 0.940 |
| | | $\alpha_2^p$ | 0.051 | 0.538 | 0.045 | 0.960 |
| | | $\alpha_3^p$ | -0.010 | 1.161 | 0.041 | 0.970 |
| | | $\alpha_4^p$ | 0.101 | 0.531 | 0.045 | 0.960 |
| | | $\alpha_5^p$ | -0.640 | 7.710 | 0.098 | 0.980 |
| | | $\sigma_f^2$ | 0.123 | 0.138 | 0.014 | 1.000 |
| | | $\sigma_\phi^2$ | 0.039 | 0.142 | 0.024 | 1.000 |
| | | $\beta^f$ | -0.040 | 0.268 | 0.043 | 0.960 |
| | | $\beta^\phi$ | 0.045 | 0.490 | 0.059 | 0.960 |
| | | $\rho^f$ | 0.120 | 0.032 | 0.042 | 1.000 |
| | | $\rho^\phi$ | 0.162 | 0.081 | 0.050 | 0.980 |
| | | $\alpha_1^f$ | 0.046 | 0.054 | 0.067 | 0.870 |
| | | $\alpha_2^f$ | -0.003 | 0.048 | 0.043 | 0.920 |
| | | $\alpha_3^f$ | 0.007 | 0.047 | 0.044 | 0.970 |
| | | $\alpha_4^f$ | 0.006 | 0.046 | 0.045 | 0.960 |
| | | $\alpha_5^f$ | 0.027 | 0.086 | 0.086 | 0.990 |
| | | $\alpha_1^\phi$ | 0.051 | 0.291 | 0.064 | 0.940 |
| | | $\alpha_2^\phi$ | -0.118 | 0.820 | 0.067 | 0.930 |
| | | $\alpha_3^\phi$ | 0.563 | 5.424 | 0.062 | 0.920 |
| | | $\alpha_4^\phi$ | -0.075 | 0.849 | 0.065 | 0.970 |
| | | $\alpha_5^\phi$ | 0.122 | 2.184 | 0.144 | 0.970 |
| Spatially constant | STM:High | $\alpha_1^p$ | -0.148 | 1.608 | 0.065 | 0.910 |
| | | $\alpha_2^p$ | 0.146 | 0.764 | 0.056 | 0.910 |
| | | $\alpha_3^p$ | 0.061 | 0.922 | 0.047 | 0.930 |
| | | $\alpha_4^p$ | -0.137 | 0.621 | 0.057 | 0.940 |
| | | $\alpha_5^p$ | -0.275 | 5.000 | 0.137 | 0.930 |
| | | $\sigma_f^2$ | 0.145 | 0.157 | 0.017 | 0.960 |
| | | $\sigma_\phi^2$ | 0.039 | 0.157 | 0.025 | 1.000 |
| | | $\beta^f$ | 0.003 | 0.389 | 0.060 | 0.930 |
| | | $\beta^\phi$ | 0.016 | 0.608 | 0.071 | 0.920 |
| | | $\rho^f$ | 0.115 | 0.031 | 0.040 | 1.000 |
| | | $\rho^\phi$ | 0.170 | 0.077 | 0.052 | 1.000 |
| | | $\alpha_1^f$ | 0.088 | 0.068 | 0.111 | 0.820 |
| | | $\alpha_2^f$ | -0.004 | 0.062 | 0.054 | 0.930 |
| | | $\alpha_3^f$ | 0.006 | 0.065 | 0.055 | 0.930 |
| | | $\alpha_4^f$ | 0.002 | 0.060 | 0.058 | 0.950 |
| | | $\alpha_5^f$ | 0.025 | 0.127 | 0.108 | 0.940 |
| | | $\alpha_1^\phi$ | 0.066 | 0.452 | 0.067 | 0.930 |
| | | $\alpha_2^\phi$ | 0.113 | 0.764 | 0.074 | 0.940 |
| | | $\alpha_3^\phi$ | -0.138 | 1.465 | 0.076 | 0.930 |
| | | $\alpha_4^\phi$ | -0.053 | 1.319 | 0.082 | 0.930 |
| | | $\alpha_5^\phi$ | 0.150 | 2.608 | 0.180 | 0.940 |

Table S4: Mean relative bias (RB), Monte Carlo standard error of RB (MCSE(RB)), root mean square error (RMSE) and 95% posterior credible interval coverage (Cov) of occasion-specific baseline recapture probability ($\alpha_t^p$), regression effects ($\beta^f, \beta^\phi$), NNGP parameters ($\sigma_f^2$, $\rho^f$, $\sigma_\phi^2, \rho^\phi$), occasion-specific baseline survival and recruitment ($\alpha_t^\phi, \alpha_t^f$) for data integration cases: perfect data alignment (PA), spatial data misalignment (SM), spatial data misalignment with low temporal data misalignment (STM:Low) and spatial data misalignment with high temporal data misalignment (STM:High) given a spatially constant covariate effect on survival and recruitment.

| Covariate effect | Case | Parameter | RB | MCSE(RB) | RMSE | Cov |
|---|---|---|---|---|---|---|
| Spatially varying | STM:Low | $\alpha_1^p$ | 0.122 | 0.734 | 0.052 | 0.950 |
| | | $\alpha_2^p$ | -0.011 | 0.354 | 0.042 | 0.950 |
| | | $\alpha_3^p$ | 0.005 | 0.896 | 0.036 | 0.990 |
| | | $\alpha_4^p$ | -0.015 | 0.559 | 0.047 | 0.930 |
| | | $\alpha_5^p$ | 0.187 | 1.825 | 0.071 | 0.980 |
| | | $\sigma_f^2$ | 0.122 | 0.127 | 0.014 | 1.000 |
| | | $\sigma_\phi^2$ | 0.024 | 0.136 | 0.021 | 1.000 |
| | | $\beta_1^f$ | 0.007 | 0.191 | 0.030 | 0.950 |
| | | $\beta_2^f$ | 0.039 | 0.889 | 0.070 | 0.910 |
| | | $\beta_3^f$ | 0.065 | 0.219 | 0.072 | 0.930 |
| | | $\beta_4^f$ | 0.019 | 0.165 | 0.027 | 0.950 |
| | | $\beta_5^f$ | 0.006 | 0.167 | 0.026 | 0.980 |
| | | $\beta_1^\phi$ | 0.011 | 0.753 | 0.057 | 0.880 |
| | | $\beta_2^\phi$ | 0.012 | 1.369 | 0.107 | 0.900 |
| | | $\beta_3^\phi$ | -0.002 | 0.253 | 0.096 | 0.920 |
| | | $\beta_4^\phi$ | 0.011 | 0.278 | 0.046 | 0.960 |
| | | $\beta_5^\phi$ | 0.005 | 0.106 | 0.043 | 0.980 |
| | | $\rho^f$ | 0.121 | 0.029 | 0.042 | 1.000 |
| | | $\rho^\phi$ | 0.153 | 0.069 | 0.047 | 1.000 |
| | | $\alpha_1^f$ | 0.031 | 0.039 | 0.044 | 0.910 |
| | | $\alpha_2^f$ | 0.011 | 0.038 | 0.036 | 0.930 |
| | | $\alpha_3^f$ | 0.012 | 0.035 | 0.031 | 0.960 |
| | | $\alpha_4^f$ | 0.003 | 0.044 | 0.041 | 0.920 |
| | | $\alpha_5^f$ | 0.010 | 0.048 | 0.044 | 0.990 |
| | | $\alpha_1^\phi$ | -0.079 | 0.724 | 0.055 | 0.930 |
| | | $\alpha_2^\phi$ | -0.035 | 0.722 | 0.057 | 0.940 |
| | | $\alpha_3^\phi$ | 0.124 | 0.626 | 0.051 | 0.930 |
| | | $\alpha_4^\phi$ | -0.035 | 0.758 | 0.071 | 0.910 |
| | | $\alpha_5^\phi$ | 0.090 | 0.659 | 0.084 | 0.990 |
| Spatially varying | STM:High | $\alpha_1^p$ | 0.054 | 1.024 | 0.055 | 0.970 |
| | | $\alpha_2^p$ | 0.058 | 0.415 | 0.050 | 0.960 |
| | | $\alpha_3^p$ | -0.125 | 1.376 | 0.050 | 0.950 |
| | | $\alpha_4^p$ | -0.047 | 0.677 | 0.055 | 0.940 |
| | | $\alpha_5^p$ | 0.004 | 3.268 | 0.084 | 0.970 |
| | | $\sigma_f^2$ | 0.141 | 0.132 | 0.016 | 1.000 |
| | | $\sigma_\phi^2$ | 0.046 | 0.139 | 0.023 | 1.000 |
| | | $\beta_1^f$ | -0.019 | 0.277 | 0.045 | 0.940 |
| | | $\beta_2^f$ | 0.120 | 0.933 | 0.069 | 0.880 |
| | | $\beta_3^f$ | 0.085 | 0.228 | 0.078 | 0.910 |
| | | $\beta_4^f$ | 0.008 | 0.214 | 0.033 | 0.930 |
| | | $\beta_5^f$ | 0.021 | 0.251 | 0.038 | 0.920 |
| | | $\beta_1^\phi$ | 0.007 | 0.768 | 0.061 | 0.960 |
| | | $\beta_2^\phi$ | 0.109 | 1.352 | 0.107 | 0.840 |
| | | $\beta_3^\phi$ | -0.003 | 0.279 | 0.111 | 0.890 |
| | | $\beta_4^\phi$ | 0.039 | 0.325 | 0.055 | 0.960 |
| | | $\beta_5^\phi$ | 0.004 | 0.114 | 0.046 | 0.980 |
| | | $\rho^f$ | 0.116 | 0.032 | 0.041 | 1.000 |
| | | $\rho^\phi$ | 0.140 | 0.073 | 0.042 | 1.000 |
| | | $\alpha_1^f$ | 0.068 | 0.046 | 0.082 | 0.800 |
| | | $\alpha_2^f$ | 0.011 | 0.046 | 0.040 | 0.950 |
| | | $\alpha_3^f$ | 0.020 | 0.052 | 0.047 | 0.940 |
| | | $\alpha_4^f$ | 0.022 | 0.058 | 0.059 | 0.920 |
| | | $\alpha_5^f$ | 0.028 | 0.069 | 0.067 | 0.940 |
| | | $\alpha_1^\phi$ | -0.020 | 0.296 | 0.056 | 0.940 |
| | | $\alpha_2^\phi$ | 0.050 | 0.458 | 0.060 | 0.950 |
| | | $\alpha_3^\phi$ | 0.695 | 6.039 | 0.070 | 0.940 |
| | | $\alpha_4^\phi$ | 0.212 | 1.431 | 0.079 | 0.940 |
| | | $\alpha_5^\phi$ | 0.145 | 1.300 | 0.115 | 0.960 |

Table S5: Mean relative bias (RB), Monte Carlo standard error of RB (MCSE(RB)), root mean square error (RMSE) and 95% posterior credible interval coverage (Cov) of occasion-specific baseline recapture probability ($\alpha_t^p$), regression effects ($\boldsymbol{\beta}^\phi, \boldsymbol{\beta}^f$), NNGP parameters ($\sigma^\nu$, $\rho^\nu$, $\sigma_\phi^2, \rho^\phi$), occasion-specific baseline survival and recruitment($\alpha_t^\phi, \alpha_t^f$) for data integration cases: spatial data misalignment with low temporal data misalignment (STM:Low) and spatial data misalignment with high temporal data misalignment (STM:High) given a spatially varying covariate effect on survival and recruitment.

| Covariate effect | Case | Parameter | RB($T=1$) | RB($T=2$) | RB($T=3$) | RB($T=4$) | RB($T=5$) | RB($T=6$) |
|---|---|---|---|---|---|---|---|---|
| Spatially constant | PA | Survival | 0.002 | 0.002 | 0.002 | 0.003 | 0.007 | |
| | | Recruitment | -0.017 | 0.010 | 0.008 | 0.015 | -0.002 | |
| | | Abundance | 0.010 | 0.002 | 0.003 | 0.003 | 0.004 | 0.006 |
| | | Growth rate | -0.000 | 0.006 | 0.005 | 0.007 | 0.007 | |
| Spatially constant | SM | Survival | 0.007 | 0.007 | 0.006 | 0.009 | 0.012 | |
| | | Recruitment | -0.015 | 0.006 | 0.009 | 0.012 | -0.002 | |
| | | Abundance | 0.010 | 0.002 | 0.003 | 0.002 | 0.004 | 0.007 |
| | | Growth rate | 0.000 | 0.006 | 0.005 | 0.007 | 0.007 | |
| Spatially constant | STM:Low | Survival | 0.008 | 0.006 | 0.008 | 0.008 | 0.018 | |
| | | Recruitment | -0.034 | 0.022 | 0.012 | 0.009 | -0.009 | |
| | | Abundance | 0.013 | 0.001 | 0.004 | 0.004 | 0.004 | 0.010 |
| | | Growth rate | -0.003 | 0.009 | 0.006 | 0.006 | 0.010 | |
| Spatially constant | STM:High | Survival | 0.006 | 0.010 | 0.009 | 0.010 | 0.015 | |
| | | Recruitment | -0.077 | 0.025 | 0.016 | 0.017 | 0.012 | |
| | | Abundance | 0.028 | -0.002 | 0.003 | 0.006 | 0.013 | 0.021 |
| | | Growth rate | -0.017 | 0.014 | 0.011 | 0.011 | 0.010 | |
| Spatially varying | STM:Low | Survival | 0.003 | 0.011 | 0.013 | 0.007 | 0.010 | |
| | | Recruitment | -0.014 | 0.004 | 0.005 | 0.014 | 0.009 | |
| | | Abundance | 0.014 | 0.002 | 0.004 | 0.004 | 0.006 | 0.011 |
| | | Growth rate | -0.002 | 0.008 | 0.006 | 0.008 | 0.009 | |
| Spatially varying | STM:High | Survival | 0.004 | 0.012 | 0.015 | 0.015 | 0.024 | |
| | | Recruitment | -0.054 | 0.009 | 0.002 | -0.003 | -0.008 | |
| | | Abundance | 0.027 | 0.001 | 0.005 | 0.006 | 0.011 | 0.022 |
| | | Growth rate | -0.012 | 0.012 | 0.009 | 0.010 | 0.014 | |

Table S6: Average mean relative bias (RB) across locations for each occasion of survival, recruitment, abundance and growth rate for data integration cases: perfect data alignment (PA), spatial data misalignment (SM), spatial data misalignment with low temporal data misalignment (STM:Low) and spatial data misalignment with high temporal data misalignment (STM:High) with spatially constant and spatially varying covariate effects on survival and recruitment.

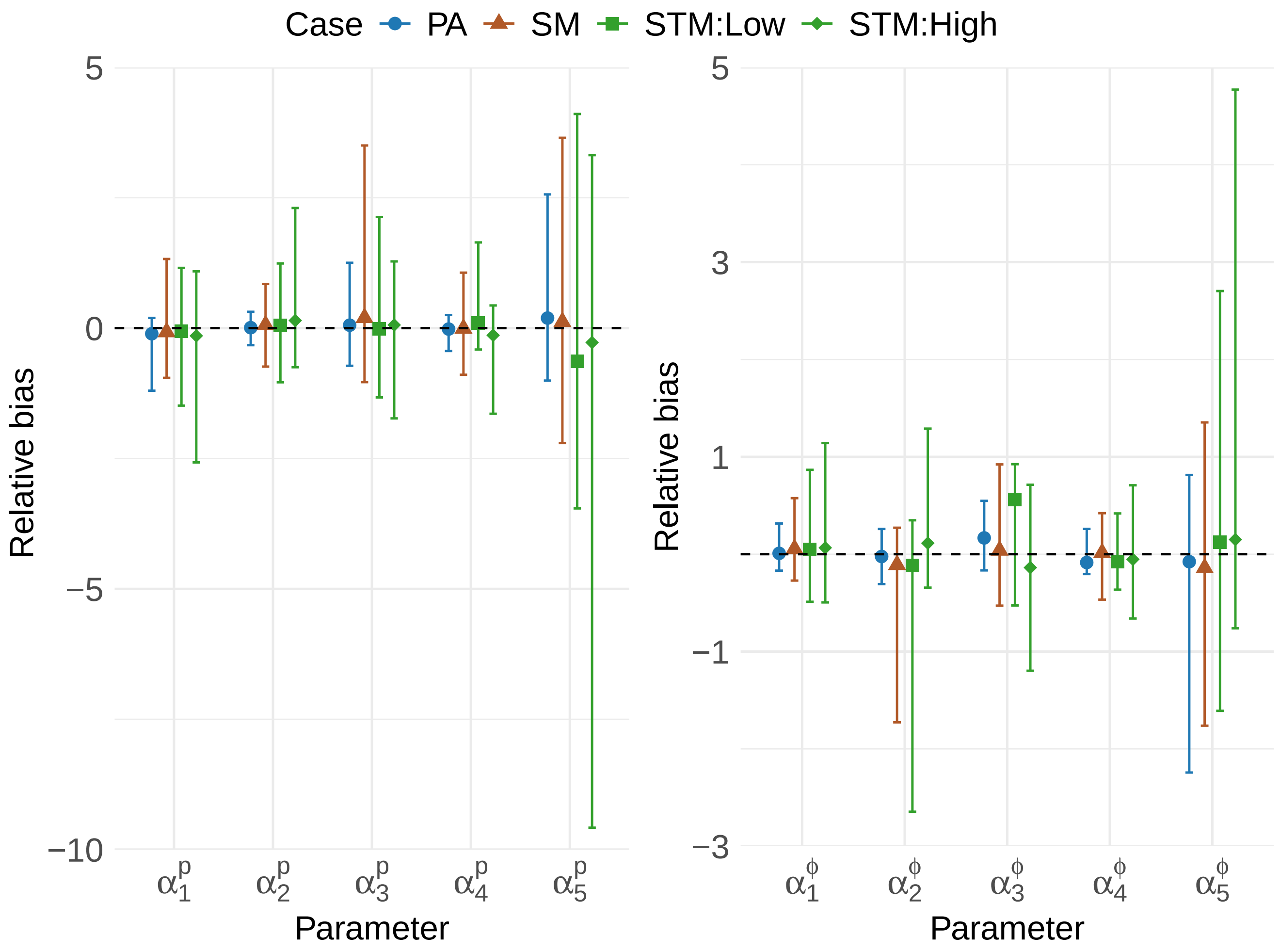


Figure S1: 95% quantile error bars of relative bias (RB) of occasion-specific baseline recapture ($\alpha_t^p$) and survival ($\alpha_t^\phi$) probabilities for data integration cases: perfect data alignment (PA), spatial data misalignment (SM), spatial data misalignment with low temporal data misalignment (STM:Low) and spatial data misalignment with high temporal data misalignment (STM:High) given a spatially constant covariate effect on survival and recruitment.

| Covariate effect | Case | Parameter | $T = 1$ | Mean RB | 2.5% RB | 97.5% RB | MAPE | RMSE | Cov |
|---|---|---|---|---|---|---|---|---|---|
| Spatially constant | PA | Survival | 1 | 0.005 | -0.106 | 0.143 | 4.961 | 0.038 | 0.958 |
| | | | 2 | 0.005 | -0.103 | 0.141 | 4.905 | 0.038 | 0.958 |
| | | | 3 | 0.005 | -0.106 | 0.144 | 4.961 | 0.038 | 0.959 |
| | | | 4 | 0.005 | -0.097 | 0.134 | 4.636 | 0.037 | 0.958 |
| | | | 5 | 0.010 | -0.094 | 0.142 | 5.141 | 0.041 | 0.961 |
| | | Recruitment | 1 | -0.010 | -0.220 | 0.248 | 10.187 | 0.047 | 0.959 |
| | | | 2 | 0.016 | -0.199 | 0.279 | 10.229 | 0.045 | 0.964 |
| | | | 3 | 0.015 | -0.201 | 0.277 | 10.201 | 0.046 | 0.963 |
| | | | 4 | 0.022 | -0.195 | 0.286 | 10.446 | 0.043 | 0.964 |
| | | | 5 | 0.005 | -0.208 | 0.264 | 10.758 | 0.046 | 0.969 |
| | SM | Survival | 1 | 0.009 | -0.116 | 0.172 | 6.039 | 0.046 | 0.955 |
| | | | 2 | 0.009 | -0.115 | 0.168 | 5.937 | 0.046 | 0.956 |
| | | | 3 | 0.008 | -0.116 | 0.171 | 6.069 | 0.046 | 0.954 |
| | | | 4 | 0.011 | -0.105 | 0.165 | 5.704 | 0.045 | 0.953 |
| | | | 5 | 0.014 | -0.105 | 0.167 | 6.739 | 0.051 | 0.949 |
| | | Recruitment | 1 | -0.008 | -0.256 | 0.311 | 12.342 | 0.056 | 0.955 |
| | | | 2 | 0.014 | -0.239 | 0.336 | 12.566 | 0.054 | 0.959 |
| | | | 3 | 0.017 | -0.236 | 0.339 | 12.523 | 0.055 | 0.959 |
| | | | 4 | 0.020 | -0.234 | 0.345 | 12.573 | 0.051 | 0.962 |
| | | | 5 | 0.006 | -0.245 | 0.328 | 13.606 | 0.056 | 0.957 |
| | STM:Low | Survival | 1 | 0.010 | -0.114 | 0.176 | 6.146 | 0.047 | 0.958 |
| | | | 2 | 0.008 | -0.115 | 0.171 | 6.130 | 0.047 | 0.953 |
| | | | 3 | 0.010 | -0.114 | 0.178 | 6.197 | 0.047 | 0.957 |
| | | | 4 | 0.009 | -0.107 | 0.165 | 5.697 | 0.045 | 0.961 |
| | | | 5 | 0.020 | -0.097 | 0.180 | 7.077 | 0.054 | 0.950 |
| | | Recruitment | 1 | -0.026 | -0.265 | 0.278 | 12.671 | 0.058 | 0.951 |
| | | | 2 | 0.030 | -0.225 | 0.353 | 12.764 | 0.054 | 0.964 |
| | | | 3 | 0.020 | -0.233 | 0.338 | 12.621 | 0.055 | 0.962 |
| | | | 4 | 0.017 | -0.232 | 0.335 | 12.674 | 0.051 | 0.966 |
| | | | 5 | -0.000 | -0.248 | 0.315 | 14.366 | 0.058 | 0.959 |
| | STM:High | Survival | 1 | 0.008 | -0.118 | 0.177 | 6.287 | 0.048 | 0.958 |
| | | | 2 | 0.011 | -0.114 | 0.177 | 6.311 | 0.048 | 0.951 |
| | | | 3 | 0.011 | -0.114 | 0.178 | 6.407 | 0.049 | 0.954 |
| | | | 4 | 0.012 | -0.107 | 0.170 | 6.080 | 0.048 | 0.953 |
| | | | 5 | 0.018 | -0.103 | 0.182 | 8.128 | 0.060 | 0.949 |
| | | Recruitment | 1 | -0.070 | -0.306 | 0.234 | 14.270 | 0.065 | 0.934 |
| | | | 2 | 0.032 | -0.231 | 0.370 | 13.809 | 0.057 | 0.963 |
| | | | 3 | 0.024 | -0.235 | 0.357 | 13.329 | 0.058 | 0.966 |
| | | | 4 | 0.026 | -0.232 | 0.364 | 13.745 | 0.054 | 0.968 |
| | | | 5 | 0.018 | -0.241 | 0.357 | 16.840 | 0.066 | 0.958 |
| Spatially varying | STM:Low | Survival | 1 | 0.005 | -0.152 | 0.219 | 6.730 | 0.045 | 0.982 |
| | | | 2 | 0.013 | -0.140 | 0.222 | 6.572 | 0.045 | 0.983 |
| | | | 3 | 0.015 | -0.142 | 0.232 | 6.740 | 0.045 | 0.983 |
| | | | 4 | 0.009 | -0.140 | 0.215 | 6.564 | 0.045 | 0.980 |
| | | | 5 | 0.012 | -0.136 | 0.212 | 6.764 | 0.047 | 0.981 |
| | | Recruitment | 1 | 0.001 | -0.257 | 0.329 | 12.482 | 0.064 | 0.985 |
| | | | 2 | 0.020 | -0.245 | 0.351 | 12.746 | 0.064 | 0.988 |
| | | | 3 | 0.021 | -0.240 | 0.355 | 12.483 | 0.064 | 0.988 |
| | | | 4 | 0.029 | -0.240 | 0.368 | 13.110 | 0.061 | 0.984 |
| | | | 5 | 0.024 | -0.241 | 0.358 | 13.043 | 0.061 | 0.985 |
| | STM:High | Survival | 1 | 0.006 | -0.157 | 0.224 | 6.937 | 0.047 | 0.982 |
| | | | 2 | 0.012 | -0.151 | 0.223 | 6.825 | 0.047 | 0.983 |
| | | | 3 | 0.017 | -0.147 | 0.239 | 7.215 | 0.048 | 0.981 |
| | | | 4 | 0.015 | -0.142 | 0.226 | 6.747 | 0.047 | 0.981 |
| | | | 5 | 0.026 | -0.126 | 0.243 | 7.382 | 0.051 | 0.980 |
| | | Recruitment | 1 | -0.038 | -0.299 | 0.293 | 13.485 | 0.069 | 0.975 |
| | | | 2 | 0.025 | -0.254 | 0.378 | 13.620 | 0.066 | 0.986 |
| | | | 3 | 0.016 | -0.256 | 0.366 | 13.548 | 0.068 | 0.984 |
| | | | 4 | 0.015 | -0.262 | 0.366 | 13.934 | 0.064 | 0.980 |
| | | | 5 | 0.009 | -0.265 | 0.351 | 14.258 | 0.066 | 0.984 |

Table S7: Mean of the mean relative bias (RB), mean 2.5% RB, mean 97.5% RB, average mean absolute percentage error (MAPE), mean root mean square error (RMSE) and mean 95% posterior credible interval coverage (Cov) across locations for each occasion of survival and recruitment predictions for data integration cases: perfect data alignment (PA), spatial data misalignment (SM), spatial data misalignment with low temporal data misalignment (STM:Low) and spatial data misalignment with high temporal data misalignment (STM:High) given spatially constant and spatially varying covariate effects respectively.

### S2 Case study: Gray Catbird

#### S2.1 Cormack-Jolly-Seber model to account for Transience with confirmation of residency status

At Monitoring Avian Productivity and Survivorship program (MAPS) sampling locations (hereafter, locations) and often in other constant effort mist surveys, an individual captured can either be a resident, that is, it belongs to the local breeding pool, or a transient. Transient individuals are individuals who merely use a location as a stopover during migration or en route during the settlement process. The presence of transient birds in the capture-recapture data leads to a negative bias in the estimates of apparent survival as by definition transient individuals have an apparent survival probability of zero (Pradel et al., 1997).

We use the Cormack-Jolly-Seber (CJS) model to filter out transient individuals and to obtain inferences on the apparent survival of resident individuals using the CR data. For the CJS model, the solution to account for transients is known (Pradel et al., 1997). For computational reasons, we employ the m-array formulation of the CJS model that accounts for transcience. In particular, we use the extension that allows confirmation of residency status within the first capture-occasion.

For $i = 1, \ldots, M_{CR}$ locations, capture histories are summarized into two m-arrays:

1. $\boldsymbol{F^m}$ - summarizing the capture histories of resident and transient individuals up to the second capture (including individuals only caught once.) $F^m_{i,j,t}$ denotes the number of individuals captured for the 1st time on occasion $j$ and then first recaptured at occasion $t$ $(j, t = 1, \ldots, T, t \geq j)$ at location $i$. $F_{i,j,j}$ denotes the number of individuals recaptured at location $i$ within the same sampling occasion as their first capture. These within-occasion recaptures provide information for confirming residency status. We assume that Catbirds captured at least twice during the breeding season, with recaptures occurring at least seven days apart are residents. $F^m_{i,j,T+1}$ represents the number of individuals captured for the first time at occasion $j$ location $i$ and never captured again at location $i$.
2. $\boldsymbol{R^m}$ - summarizing residents from the second capture onward (second capture when an individual is a confirmed resident). $R^m_{i,j,t}$ denotes the number of individuals (i.e. confirmed residents) at occasion $j$ and next recaptured at occasion $t$ $(j = 1, \ldots, T-1, t = 2, \ldots, T, t > j)$ at location $i$ and $R^m_{i,j,T+1}$ denotes the number of individuals recaptured at occasion $j$ location $i$ and never recaptured again.

The full likelihood of the CJS transience model is the product of the likelihoods of $\boldsymbol{F^m}$ and $\boldsymbol{R^m}$. We let $\boldsymbol{F^m}$ follow a multinomial-based likelihood where for each location, each row $j$ of $\boldsymbol{F^m}$ follows an independent multinomial distribution defined as

$$F^m_{i,j,1:T+1} \sim \text{Multinomial}(\sum_{t=1}^{T+1} F^m_{i,j,t}, f^m_{i,j,1:T+1})$$

where $f^m_{i,j,1:T}$ are the m-array cell probabilities representing the probability that an individual captured for the first time at occasion $j$ location $i$ is first recaptured at occasion $t$ location $i$. $f^m_{i,j,T+1}$ represents the probability that an individual captured for the first time at occasion $j$ location $i$ is never recaptured again. To define the $f^m_{i,j,1:T+1}$, let $r^F_j$ denote the probability that an animal captured for the first time on occasion $j$ is a resident (residency probability within newly captured individuals), $\psi_j$ represent the probability of the confirmation of residency status for the newly captured resident, $\phi_{i,t}$ denotes apparent survival of residents at location $i$ from occasion $t$ to $t+1$ and $p_{i,t}$ denotes the recapture probability of a resident individual at location $i$ and occasion $t+1$. Consequently,

$$f^m_{i,j,j} = r^F_j \psi_j$$

$$f^m_{i,j,t} = r^F_j (1 - \psi_j) \phi_{i,t-1} p_{i,t-1} \prod_{l=j}^{t-2} \phi_{i,l} (1 - p_{i,l}) \qquad j < t \leq T$$

$$f^m_{i,j,T+1} = 1 - \sum_{t=1}^{T} f^m_{i,j,t}$$

Thus, the likelihood of $\boldsymbol{F^m}$ is the product of the multinomial distribution defined above for each location and each occasion $j$ of $F^m_{i,j,1:T+1}$.

Likewise, we let $\boldsymbol{R^m}$ follow a multinomial-based likelihood where for each location, each row $j$ of $\boldsymbol{R^m}$ follows an independent multinomial distribution defined as

$$R^m_{i,j,2:T+1} \sim \text{Multinomial}\left(\sum_{t=2}^{T+1} R^m_{i,j,t}, r^m_{i,j,2:T+1}\right)$$

where the cell probabilities $r^m_{i,j,t}$ represents the probability that an individual recaptured at occasion $j$ location $i$ is next recaptured at occasion $t$ and $r^m_{i,j,T+1}$ represents the probability that an individual recaptured at occasion $j$ location $i$ is never recaptured again can be expressed as in the standard CJS m-array way:

$$r^m_{i,j,t} = \phi_{i,t-1} p_{i,t-1} \prod_{l=j}^{t-2} \phi_{i,l}(1 - p_{i,l}) \qquad j < t \leq T$$

$$r^m_{i,j,T+1} = 1 - \sum_{t=1}^{T} r^m_{i,j,t}$$

The likelihood of $\boldsymbol{R^m}$ is the product of the multinomial distribution defined above for each location and each occasion $j$ of $R^m_{i,j,2:T+1}$.

#### S2.2 Spatio-temporal misalignment

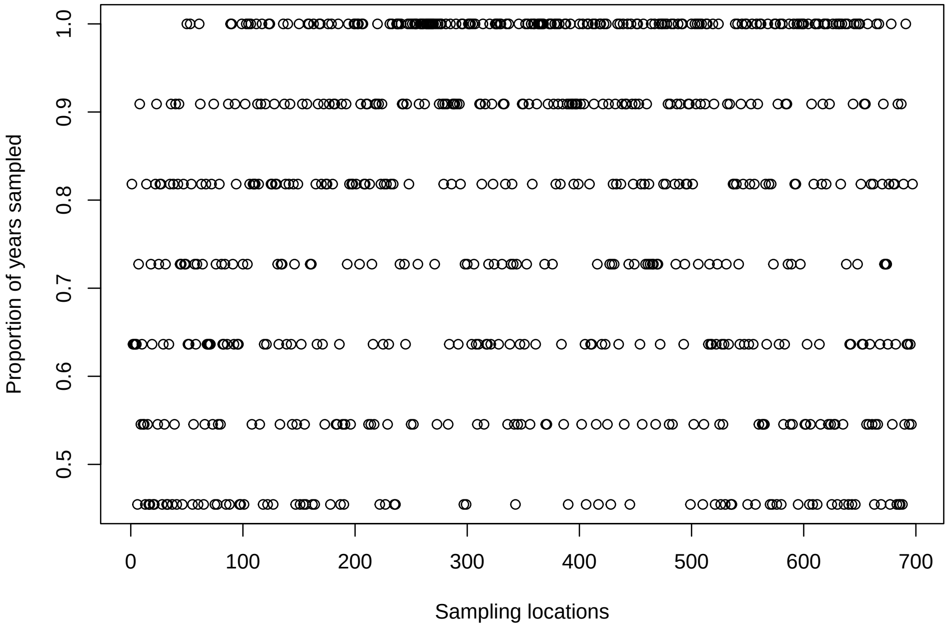


Figure S2: Proportion of years sampled at each location in the North American Breeding Bird Survey (BBS). Points represent individual sampling locations.

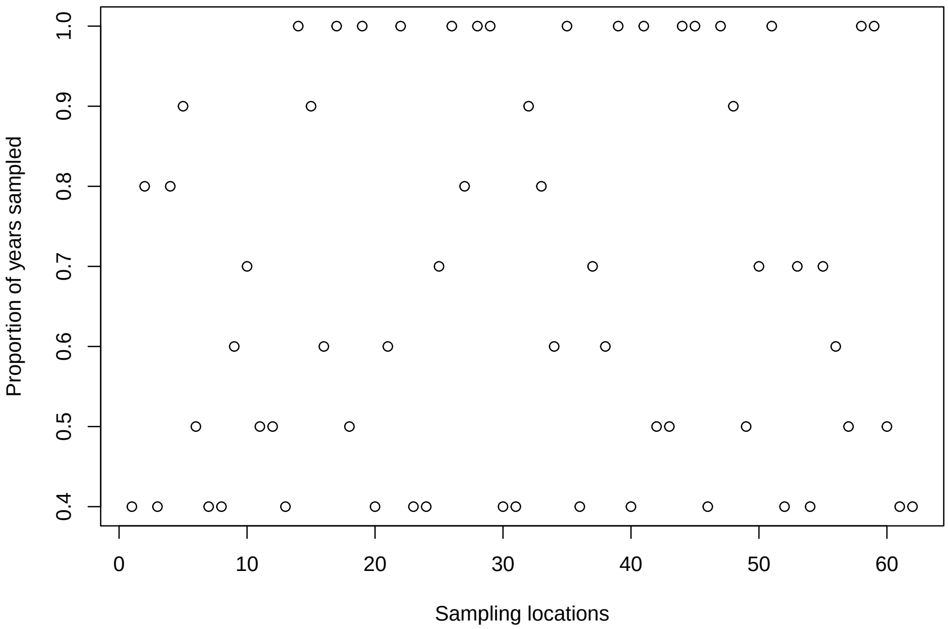


Figure S3: Proportion of years sampled at each location in the Monitoring Avian Productivity and Survivorship program (MAPS). Points represent individual sampling locations.

#### S2.3 Settings

We considered two cases of residual maximum temperature ($RTMAX$) effect on survival and recruitment during the breeding season (May-August): a spatially constant case and a spatially varying case. $RTMAX$ was defined as the residuals from a linear model of maximum temperature on an intercept and spatial coordinates. Let $\nu \in \{\phi, f\}$. For the spatially constant case:

$$g(RTMAX_{i,t}, \beta^{\nu}) = RTMAX_{i,t}\beta^{\nu}$$

For the spatially varying case, we considered an interaction function between spatial coordinates and $RTMAX$:

$$g(RTMAX_{i,t}, \beta^{\nu}) = RTMAX_{i,t}\beta_1^{\nu} + Long_i\beta_2^{\nu} + Lat_i\beta_3^{\nu} + (RTMAX_{i,t} \times Long_i)\beta_4^{\nu} + (RTMAX_{i,t} \times Lat_i)\beta_5^{\nu}$$

where $Lat_i, Long_i$ denotes the latitude and longitude of location $i$ respectively.

| Parameter | Prior |
|---|---|
| Initial abundance | Discrete Uniform (1, 100) |
| $\alpha_t^p$ | Normal (0, $\sigma^2 = 2$) |
| $r_j^F$ | Uniform(0,1) |
| $\psi_j$ | Uniform(0,1) |
| $\beta^\nu$ | Normal(0, $\sigma^2$= 2) |
| $\beta_1^\nu$ | Normal(0, $\sigma^2$= 2) |
| $\beta_2^\nu$ | Normal(0, $\sigma^2$= 2) |
| $\beta_3^\nu$ | Normal(0, $\sigma^2$= 2) |
| $\beta_4^\nu$ | Normal(0, $\sigma^2$= 2) |
| $\beta_5^\nu$ | Normal(0, $\sigma^2$= 2) |
| $\eta$ | Normal(0, $\sigma^2 = 2$) |
| $\upsilon_\ell$ | Normal(0, $\sigma^2 = 2$) |
| $\rho_f$ | Uniform($\frac{dmin}{3}$, $\frac{dmax}{3}$) |
| $\sigma_f^2$ | Inverse Gamma(3,1) |
| $\rho_\phi$ | Uniform($\frac{dmin}{3}$, $\frac{dmax}{3}$) |
| $\sigma_\phi^2$ | Inverse Gamma(3,1) |
| $\alpha_t^\phi$ | Normal (0, $\sigma^2 = 2$) |
| $\alpha_t^f$ | Normal (0, $\sigma^2 = 2$) |

Table S8: Gray Catbird prior settings of occasion-specific baseline recapture probability ($\alpha_t^p$), residency probability within newly captured individuals ($r_j^F$), probability of the confirmation of residency status for the newly captured resident ($\psi_j$), spatially constant regression effects ($\beta^\nu$), spatially varying regression effects ($\beta_{1:5}^\nu$), first time observer regression effect ($\eta$), observer random effect ($\upsilon_\ell$), Nearest Neighbor Gaussian Process parameters ($\sigma_f^2$, $\rho^f$, $\sigma_\phi^2, \rho^\phi$) of survival and recruitment, occasion-specific baseline survival ($\alpha_t^\phi$) and occasion-specific baseline recruitment ($\alpha_t^f$). $dmin, dmax$ denote minimum and maximum euclidean distance respectively.

Table S9 displays the MCMC settings used in each case. In all cases considered, parameters in Table S8 were thinned by 30 while survival, recruitment, population abundance and population growth rate where thinned by 100.

| Covariate effect | NNGP | Iterations | Burn-in | Chains |
|---|---|---|---|---|
| Spatially varying | No | 250000 | 150000 | 2 |
| Spatially constant | No | 250000 | 150000 | 2 |
| Spatially varying | Yes | 350000 | 200000 | 2 |
| Spatially constant | Yes | 350000 | 200000 | 2 |

Table S9: MCMC settings for cases considered for the different covariate effects and whether the Nearest Neighbor Gaussian Process (NNGP) was used.

#### S2.4 Residual Maximum Temperature

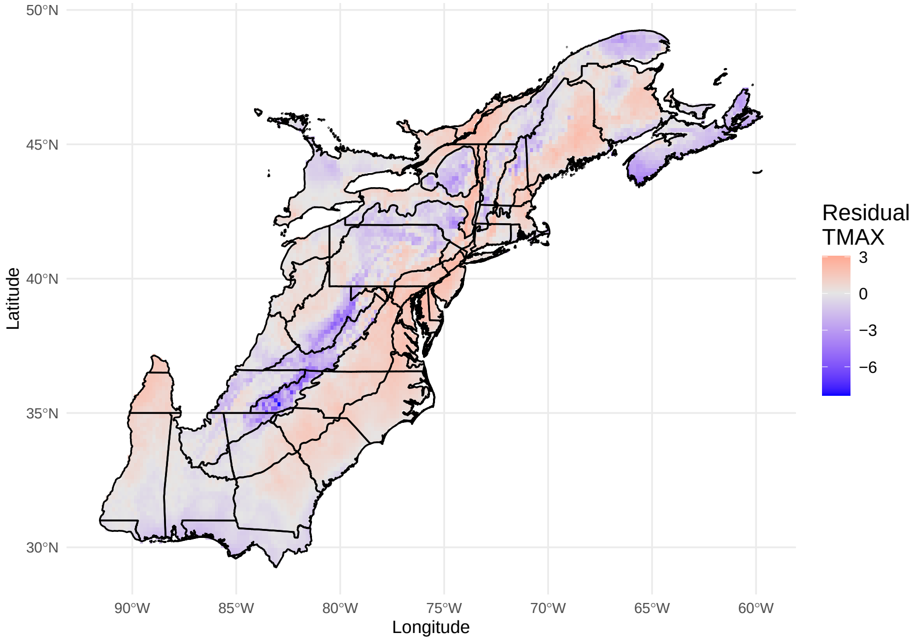


Figure S4: Spatial distribution of average residual maximum temperature ($TMAX$) on BCR 30, 29, 28, 27, 14, 13 regions across 2004-2014.

#### S2.5 Posterior Predictive loss ($PPL$)

Posterior Predictive loss ($PPL$) is a Bayesian model selection criterion that measures how well a model's predicted data ($\tilde{y}_i$), generated from its posterior predictive distribution, matches the observed ($\boldsymbol{y}$). PPL is defined as a combination of goodness-of-fit ($G$) and penalty ($P$):

$$PPL = G + P$$

$$G = \sum_{i=1}^{n} (y_i - E(\tilde{y}_i|\boldsymbol{y}))^2$$

$$P = \sum_{i=1}^{n} Var(y_i|\boldsymbol{y})$$

where $E(\tilde{y}_i|\boldsymbol{y})$ is the posterior prediction mean and $Var(y_i|\boldsymbol{y})$ is the posterior prediction variance for $\tilde{y}_i$. It balances fit and penalty by rewards models that fit well and penalizing overfitted models as $\sum_{i=1}^{n} Var(y_i|\boldsymbol{y})$ becomes larger with an increasing number of parameters. Thus, lower PPL indicates better predictive performance. For $\ell = 1, \ldots, L$ different data types, $PPL$ can be defined

$$PPL = \sum_{\ell=1}^{L} \sum_{i=1}^{n} (y_i^\ell - E(\tilde{y}_i^\ell|\boldsymbol{y}^\ell))^2 + \sum_{i=1}^{n} Var(y_i^\ell|\boldsymbol{y}^\ell)$$

where $\tilde{y}_i^\ell$ is the model's predictive data for data type $\ell$ and $\boldsymbol{y}^\ell$ is the observed data type $\ell$.

#### S2.6 Results

#### S2.7 MCMC summaries

From Table S9, residency probability and probability of confirmation residency status, components of the m-array CJS model that accounts for transience, were found to be very similar to the estimates of Ahrestani et al. (2017) and Schaub and Kéry (2022).

| Parameter | Mean | Standard deviation | 95% Posterior Credible Interval |
|---|---|---|---|
| $\alpha_1^p$ | -0.365 | 0.334 | (-1.019,0.288) |
| $\alpha_2^p$ | -0.929 | 0.303 | (-1.51,-0.325) |
| $\alpha_3^p$ | -0.638 | 0.293 | (-1.22,-0.064) |
| $\alpha_4^p$ | -0.740 | 0.317 | (-1.364,-0.127) |
| $\alpha_5^p$ | -0.917 | 0.315 | (-1.532,-0.3) |
| $\alpha_6^p$ | -0.596 | 0.321 | (-1.231,0.035) |
| $\alpha_7^p$ | -0.277 | 0.317 | (-0.896,0.352) |
| $\alpha_8^p$ | -0.910 | 0.320 | (-1.54,-0.281) |
| $\alpha_9^p$ | -0.934 | 0.341 | (-1.602,-0.254) |
| $\alpha_{10}^p$ | -0.630 | 0.867 | (-2.078,1.161) |
| $\sigma_f^2$ | 0.046 | 0.008 | (0.032,0.064) |
| $\sigma_\phi^2$ | 0.079 | 0.018 | (0.05,0.121) |
| $\beta_1^f$ | -0.055 | 0.084 | (-0.22,0.107) |
| $\beta_2^f$ | 0.074 | 0.073 | (-0.061,0.219) |
| $\beta_3^f$ | 0.499 | 0.620 | (-0.686,1.735) |
| $\beta_4^f$ | -0.404 | 0.374 | (-1.163,0.311) |
| $\beta_5^f$ | 0.116 | 0.398 | (-0.639,0.931) |
| $\beta_1^\phi$ | 0.162 | 0.151 | (-0.131,0.441) |
| $\beta_2^\phi$ | -0.153 | 0.137 | (-0.425,0.095) |
| $\beta_3^\phi$ | -0.805 | 0.989 | (-2.73,1.151) |
| $\beta_4^\phi$ | 0.966 | 0.605 | (-0.218,2.15) |
| $\beta_5^\phi$ | 0.117 | 0.708 | (-1.275,1.483) |
| $\eta$ | -0.073 | 0.026 | (-0.124,-0.022) |
| $r_1^F$ | 0.671 | 0.094 | (0.518,0.892) |
| $r_2^F$ | 0.559 | 0.063 | (0.45,0.694) |
| $r_3^F$ | 0.506 | 0.054 | (0.409,0.624) |
| $r_4^F$ | 0.615 | 0.065 | (0.502,0.755) |
| $r_5^F$ | 0.590 | 0.066 | (0.476,0.733) |
| $r_6^F$ | 0.489 | 0.055 | (0.391,0.609) |
| $r_7^F$ | 0.678 | 0.083 | (0.535,0.861) |
| $r_8^F$ | 0.557 | 0.055 | (0.461,0.674) |
| $r_9^F$ | 0.570 | 0.075 | (0.444,0.738) |
| $r_{10}^F$ | 0.715 | 0.107 | (0.522,0.942) |
| $r_{11}^F$ | 0.715 | 0.107 | (0.522,0.942) |
| $\rho_f^2$ | 939.982 | 46.427 | (817.518,987.828) |
| $\rho_\phi^2$ | 901.796 | 77.248 | (697.482,986.753) |
| $\psi_1$ | 0.273 | 0.043 | (0.193,0.36) |
| $\psi_2$ | 0.336 | 0.045 | (0.251,0.428) |
| $\psi_3$ | 0.378 | 0.047 | (0.29,0.473) |
| $\psi_4$ | 0.289 | 0.038 | (0.216,0.367) |
| $\psi_5$ | 0.360 | 0.045 | (0.276,0.452) |
| $\psi_6$ | 0.425 | 0.053 | (0.325,0.532) |
| $\psi_7$ | 0.266 | 0.040 | (0.193,0.35) |
| $\psi_8$ | 0.369 | 0.043 | (0.288,0.454) |
| $\psi_9$ | 0.345 | 0.051 | (0.251,0.45) |
| $\psi_{10}$ | 0.218 | 0.040 | (0.15,0.304) |
| $\psi_{11}$ | 0.218 | 0.040 | (0.15,0.304) |
| $\alpha_1^f$ | -0.897 | 0.211 | (-1.372,-0.571) |
| $\alpha_2^f$ | -1.226 | 0.334 | (-2.065,-0.703) |
| $\alpha_3^f$ | -0.742 | 0.154 | (-1.108,-0.484) |
| $\alpha_4^f$ | -0.941 | 0.268 | (-1.572,-0.553) |
| $\alpha_5^f$ | -0.762 | 0.198 | (-1.215,-0.472) |
| $\alpha_6^f$ | -0.783 | 0.209 | (-1.287,-0.486) |
| $\alpha_7^f$ | -0.716 | 0.149 | (-1.031,-0.461) |
| $\alpha_8^f$ | -0.922 | 0.238 | (-1.493,-0.554) |
| $\alpha_9^f$ | -1.123 | 0.389 | (-2.081,-0.65) |
| $\alpha_{10}^f$ | -0.685 | 0.487 | (-2.2,-0.22) |
| $\alpha_1^\phi$ | -0.098 | 0.339 | (-0.726,0.569) |
| $\alpha_2^\phi$ | 0.530 | 0.407 | (-0.261,1.436) |
| $\alpha_3^\phi$ | 0.257 | 0.294 | (-0.278,0.911) |
| $\alpha_4^\phi$ | 0.469 | 0.433 | (-0.254,1.432) |
| $\alpha_5^\phi$ | -0.007 | 0.355 | (-0.602,0.754) |
| $\alpha_6^\phi$ | 0.439 | 0.406 | (-0.186,1.373) |
| $\alpha_7^\phi$ | -0.170 | 0.288 | (-0.728,0.397) |
| $\alpha_8^\phi$ | 0.390 | 0.399 | (-0.292,1.286) |
| $\alpha_9^\phi$ | 0.344 | 0.476 | (-0.418,1.446) |
| $\alpha_{10}^\phi$ | -0.059 | 0.895 | (-1.205,2.441) |

Table S10: MCMC posterior summary for occasion-specific baseline recapture probability ($\alpha_t^p$), NNGP parameter ($\sigma_f^2, \sigma_\phi^2, \rho^f, \rho^\phi$), regression effects ($\boldsymbol{\beta}^f, \boldsymbol{\beta}^\phi$), first time observer effect ($\eta$), temporal residency probability ($r_j^F$), temporal probability of confirmation residency status ($\psi_j$), occasion-specific baseline recruitment ($\alpha_t^f$) and occasion-specific baseline survival ($\alpha_t^\phi$).

#### S2.8 Effective spatial range

The effective spatial range ($h_{\text{eff}}$) is the distance beyond which spatial correlation is considered negligible. Mathematically, it is as the distance where correlation falls to a small value $\epsilon$. In this work, we employ an exponential covariance function:

$$K(h) = \sigma^2 \exp(-h/\rho)$$

where $h$ is the distance between two locations. This results in the correlation function

$$\text{Corr}(h) = \exp(-h/\rho).$$

We let $\epsilon = 0.05$, thus

$$\epsilon = \exp(-h_{\text{eff}}/\rho)$$

and solving for $h_{\text{eff}}$:

$$h_{\text{eff}} \approx 3\rho$$

The effective spatial range of residual spatial autocorrrelation, which is the distance beyond which spatial autocorrelation is considered negligible, was 2705.388 km with $(2092.446, 2960.259)$ km 95% posterior credible interval for survival and 2819.947 km with $(2452.555, 2963.485)$ km 95% PCI for recruitment. Figure S5 displays a plot of the effective spatial range for survival and recruitment residual spatial autocorrrelation.

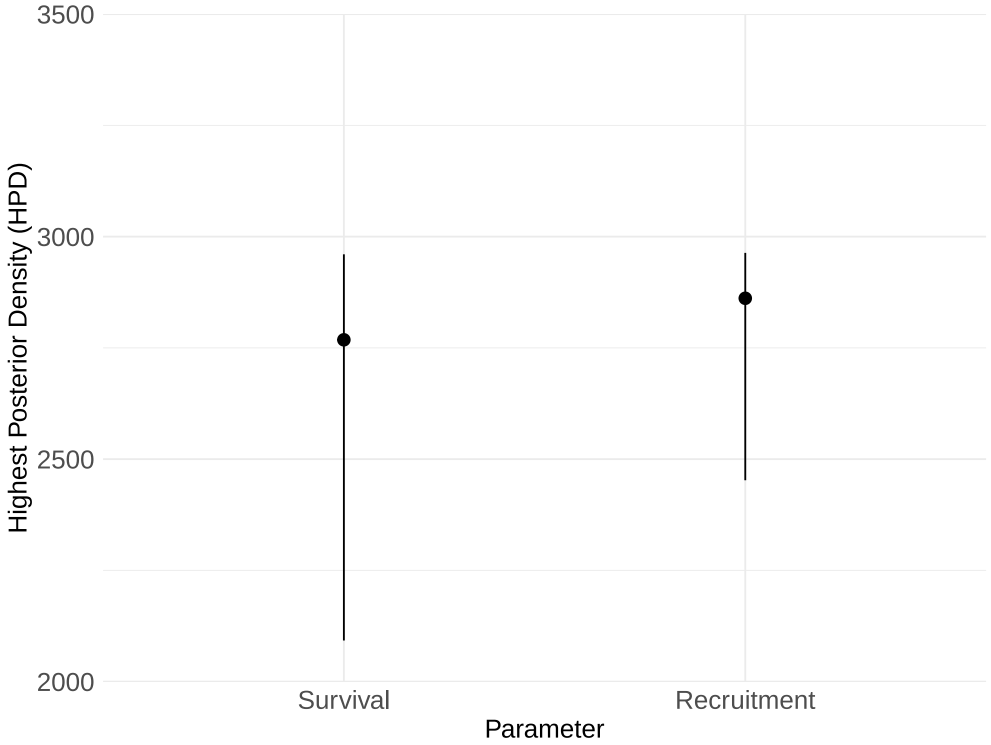


Figure S5: Effective spatial range of survival and recruitment residual spatial autocorrelation of Gray Catbirds.

#### S2.9 Prediction uncertainty

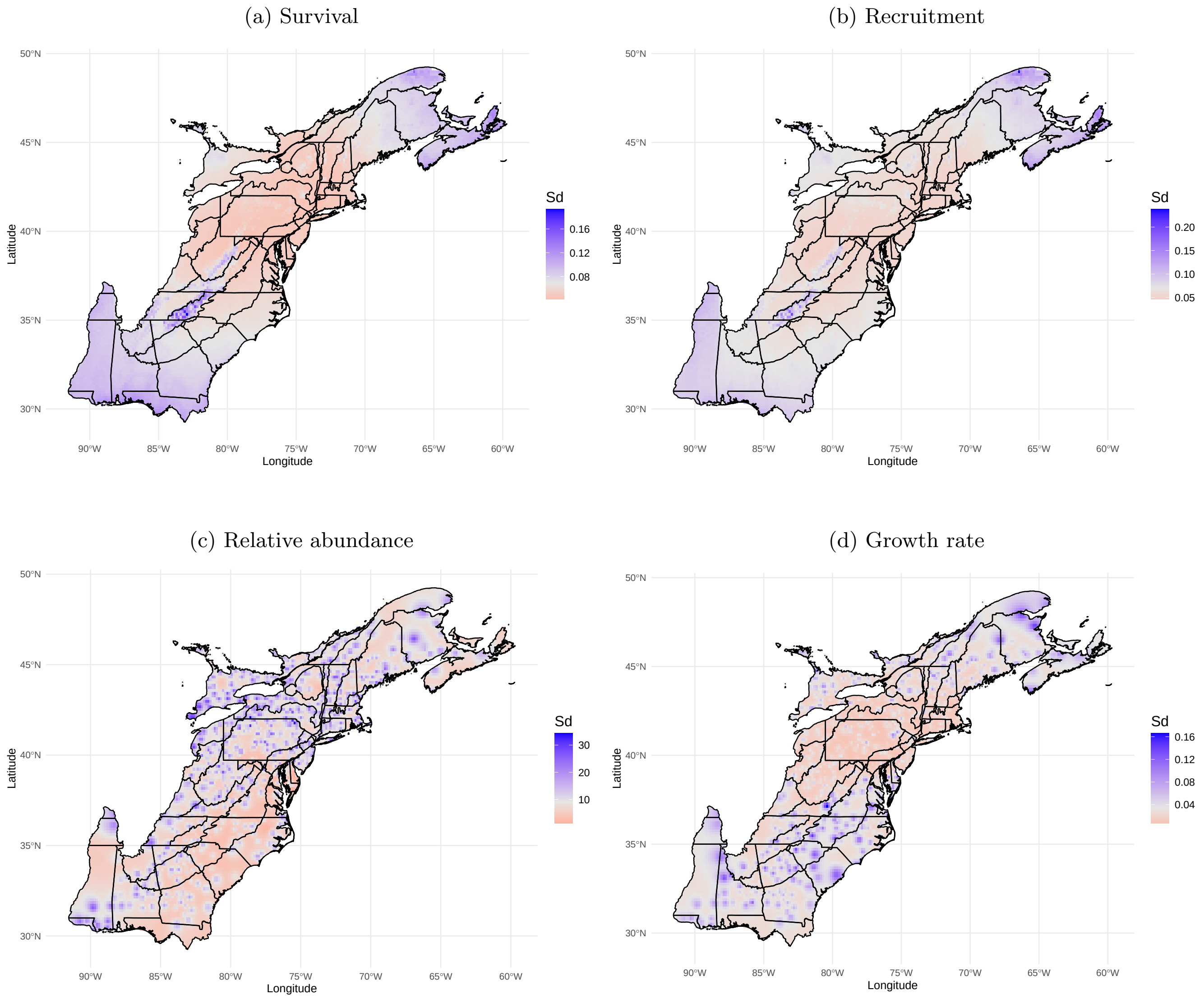


Figure S6: Maps showing the standard deviation (Sd) of spatial predictions for survival, recruitment, relative abundance and growth rate of Gray Catbirds averaged across 2004-2014.

#### S2.10 Residual Maximum Temperature Slope Uncertainty

Given we employ an interaction function between the spatial coordinates and residual maximum temperature, as described the S2.3, the slope at location $i$ can be computed as

$$\text{Slope}_i^{\nu} = \beta_1^{\nu} + Long_i\beta_4^{\nu} + Lat_i\beta_5^{\nu}.$$

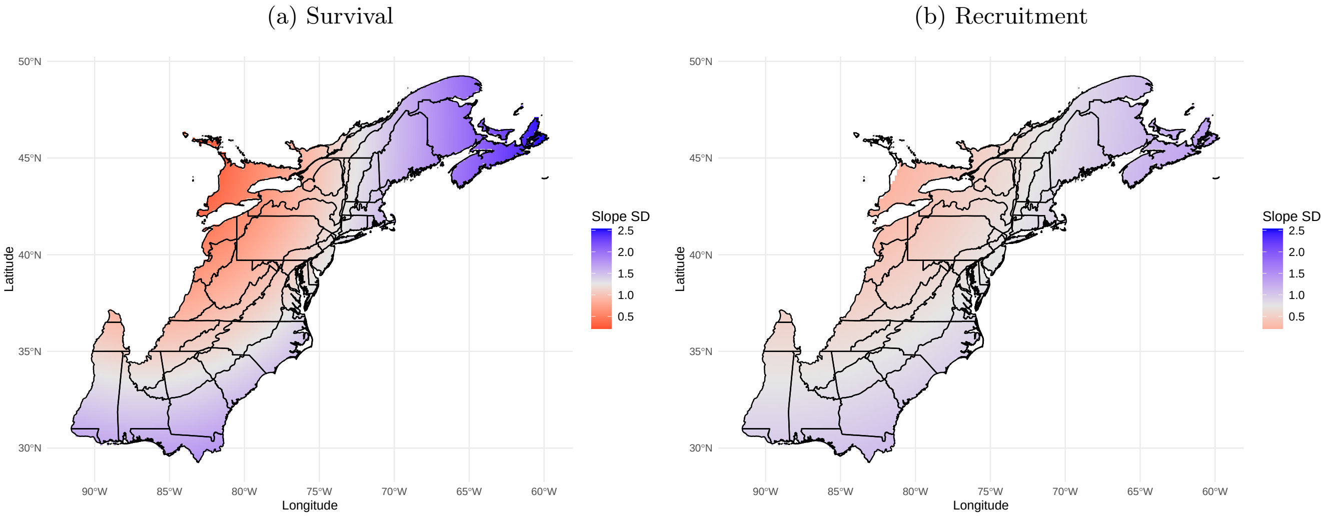


Figure S7: Spatially varying slope standard deviation of residual maximum temperature for survival and recruitment of Gray Catbirds across the eastern coast of the United States from 2004-2014.

#### S2.11 Posterior predictive checks

For $i = 1, \ldots, M$ locations, $t = 1, \ldots, T$ time periods and $l = 1, \ldots, L$ MCMC samples. For a Poisson distribution, the expected mean is the rate. For a multinomial distribution, the expected mean is $N\pi$ where $N$ is the size and $\pi$ is the cell probabilities. Throughout this section we use $BPV$ to denote Bayesian p-value and $FT$ to denote Freeman-Tukey fit statistic.

##### S2.11.1 North American Breeding Bird Survey count data

Let $C_{i,t}$ be the observed count at location $i$ time $t$, $C_{pred,i,t}$ be the predicted data at location $i$ time $t$ and $\lambda_{i,t}$ is the expected mean.

###### Spatial Grouping

Define $C_{i,.}$ and $C^l_{pred,i,.}$ as the total number of counts at location $i$ for the observed data and the prediction data during iteration $l$, respectively.

$$C_{i,.} = \sum_t C_{i,t}$$

$$C^l_{pred,i.} = \sum_t C^l_{pred,i,t}$$

$$FT^l_i = (\sqrt{C_{i,.}} - \sqrt{\lambda^l_{i,.}})^2$$

$$FT^l_{pred,i} = (\sqrt{C^l_{pred,i,.}} - \sqrt{\lambda^l_{i,.}})^2$$

Count location specific $BPV$ $(BPV^C_i)$ is computed as

$$BPV^C_i = \frac{\sum_l I(FT^l_{pred,i} > FT^l_i)}{L}$$

while count global spatial $BPV$ $(BPV^C_{Gs})$ is computed as

$$BPV^C_{Gs} = \frac{\sum_l I(\sum_i FT^l_{pred,i} > \sum_i FT^l_i)}{L}$$

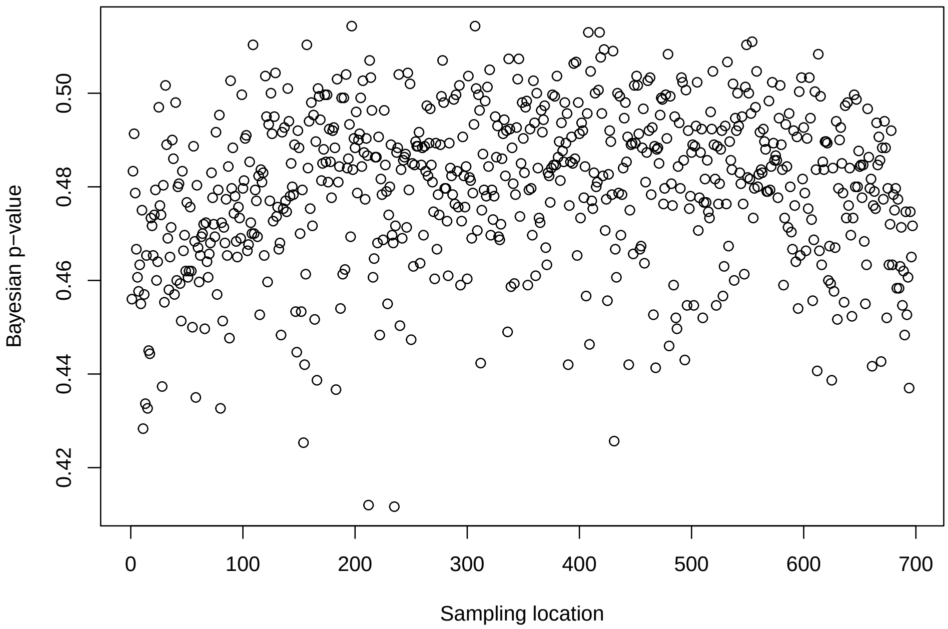


Figure S8: Sampling location specific Bayesian p-values ($BPV$) for North American Breeding Bird Survey (BBS) count data.

Global spatial $BPV = 0.567$.

**Temporal grouping**

Define $C_{.,t}$ and $C^l_{pred,.,t}$ as the total number of counts at time $t$ for the observed data and the prediction data during iteration $l$, respectively.

$$C_{.,t} = \sum_i C_{i,t}$$
$$C^l_{pred,.,t} = \sum_i C^l_{pred,i,t}$$
$$FT^l_t = (\sqrt{C_{.,t}} - \sqrt{\lambda^l_{.,t}})^2$$
$$FT^l_{pred,t} = (\sqrt{C^l_{pred,.,t}} - \sqrt{\lambda^l_{.,t}})^2$$

Count time specific BPV ($BPV^C_t$) is computed as

$$BPV^C_t = \frac{\sum_l I(FT^l_{pred,t} > FT^l_t)}{L}$$

while the count global temporal $BPV$ ($BPV^C_{Gt}$) is computed as

$$BPV^C_{Gt} = \frac{\sum_l I(\sum_t FT^l_{pred,t} > \sum_t FT^l_t)}{L}$$

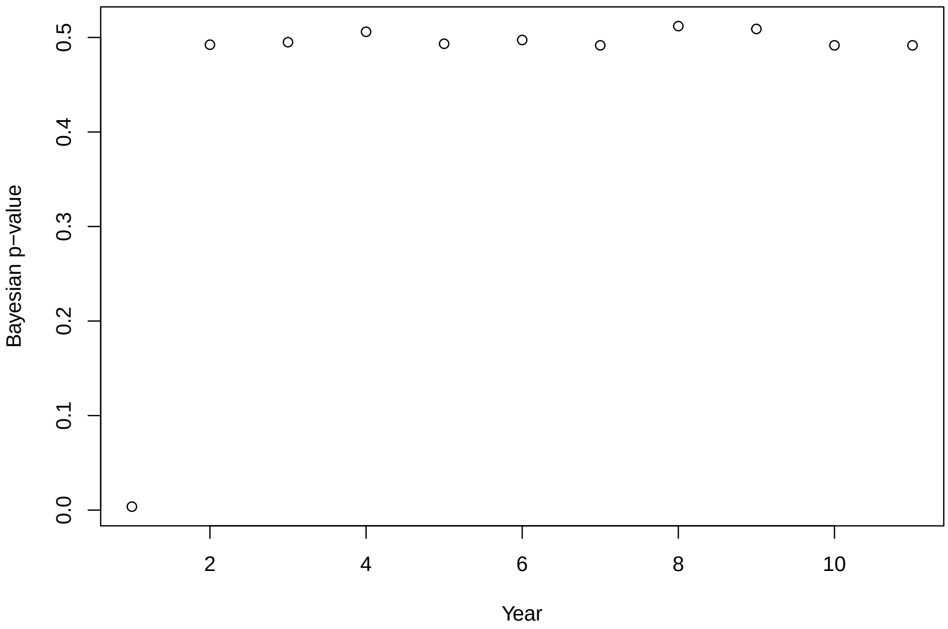


Figure S9: Year specific Bayesian p-values ($BPV$) for North American Breeding Bird Survey (BBS) count data.

Global temporal BPV = 0.0606. Global temporal BPV excluding 1st year 0.493.

##### S2.11.2 Monitoring Avian Productivity and Survivorship program (MAPS) CR data

$R_{i,1:(T-1),1:T}$ is the m-arry at location $i$ and the expected mean $\lambda_{i,1:(T-1),1:T} = N_{i,t} * \pi_{i,1:(T-1),1:T}$ where $N_{i,t}$ is the number of releases and $\pi_{i,1:(T-1),1:T}$ is the cell probabilities. Let $i = 1, \ldots, M$ locations, $t = 1, \ldots, T-1$ capture occasions and $k = 1, \ldots, T$ recapture occasions. We further denote Let $R_{pred,i,t,k}$ as the prediction data.

###### Spatial Grouping

$$R_{i,t,.} = \sum_k R_{i,t,k}$$
$$R_{pred,i,t,.} = \sum_k R_{pred,i,t,k}$$
$$FT^l_{i,t} = (\sqrt{R}_{i,t,.} - \sqrt{\lambda}^l_{i,t,})^2$$
$$FT^l_{pred,i,t} = (\sqrt{R}^l_{pred,i,t,.} - \sqrt{\lambda}^l_{i,t,.})^2$$

The CR location specific BPV ($BPV_i^{CR}$) is computed as

$$BPV_i^{CR} = \frac{\sum_l I(\sum_t FT^l_{pred,i,t} > \sum_t FT^l_{i,t})}{L}$$

and the CR global spatial $BPV$ $BPV_{Gs}^{CR}$ is computed as

$$BPV_{Gs}^{CR} = \frac{\sum_l I(\sum_t \sum_k FT^l_{pred,i,t} > \sum_t \sum_k FT^l_{i,t})}{L}$$

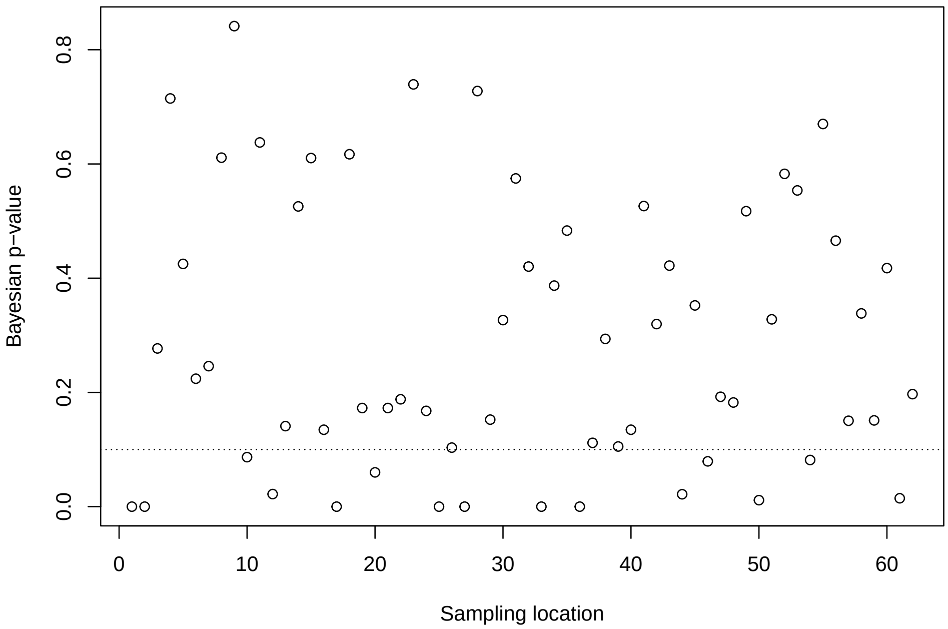


Figure S10: Samplong location specific Bayesian p-value of resident individuals from the Monitoring Avian Productivity and Survivorship (MAPS) program. Dashed horizontal line Bayesian p-value= 0.1

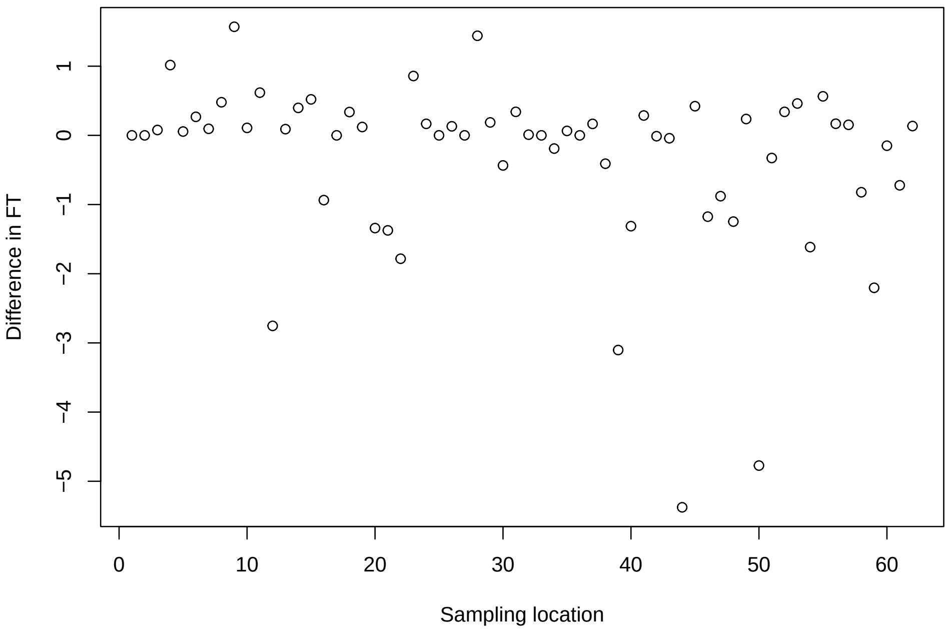


Figure S11: Sampling location specific difference in Freeman-Tukey (FT) statistics of resident individuals from the Monitoring Avian Productivity and Survivorship (MAPS) program.

CR global spatial $BPV = 0.016$. From Figure S10, 15 sampling location had $BPV > 0.1$. However, 7 of these sampling locations had 0 captures and releases of known residences. Thus, only 8 MAPS sampling locations did not seem to fit the data well. The difference in FT (Figure S11) doesn't seem to be large indicating similar fit.

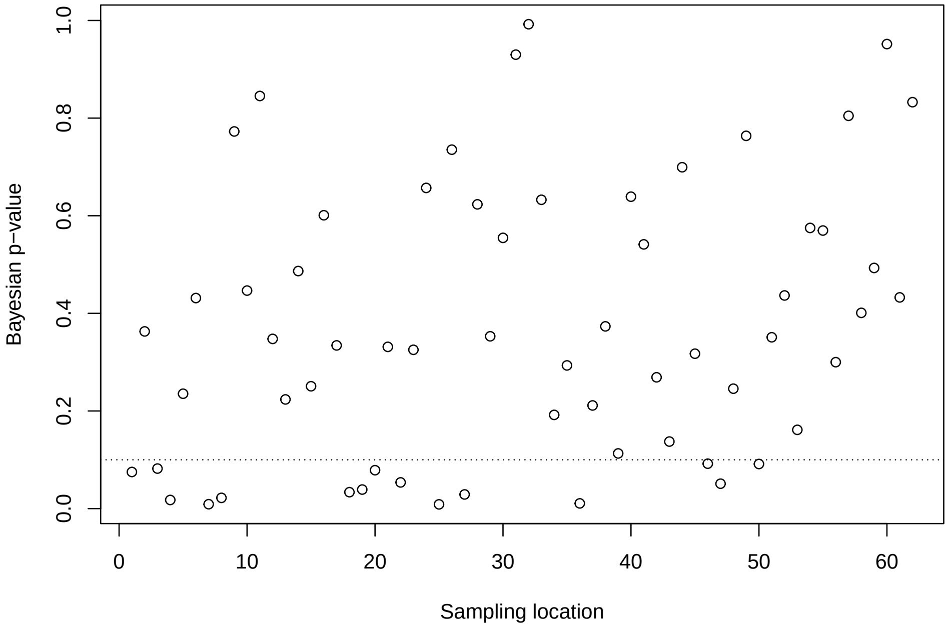


Figure S12: Sampling location specific Bayesian p-value of possible transient and resident individuals from the Monitoring Avian Productivity and Survivorship (MAPS) program. Dashed horizontal line Bayesian p-value= 0.1

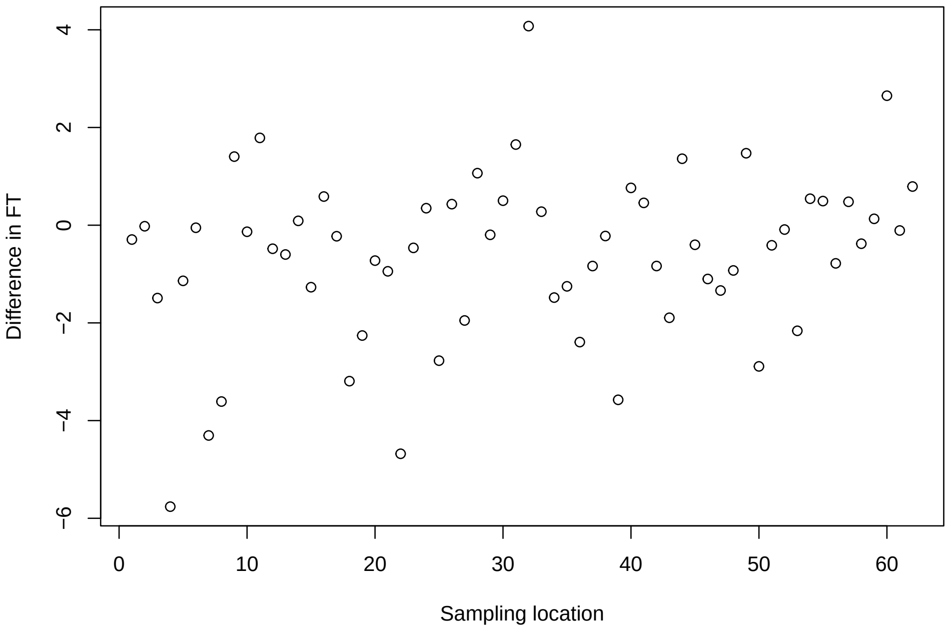


Figure S13: Sampling location specific difference in Freeman-Tukey (FT) statistics of possible transient and resident individuals from the Monitoring Avian Productivity and Survivorship (MAPS) program.

CR global spatial BPV = 0.003 for the mixture of transient and resident. From Figure S12, 14 sampling location had $BPV > 0.1$. However, as seen from Figure S13, the difference in FT statistic is not large for locations that suggest inadequate fit.

**Temporal grouping**

$$R_{.,t,k} = \sum_i R_{i,t,k}$$
$$R_{pred,.,t,k} = \sum_i R_{pred,i,t,k}$$
$$FT^l_{t,k} = (\sqrt{R}_{.,t,k} - \sqrt{\lambda}^l_{.,t,k})^2$$
$$FT^l_{pred,t,k} = (\sqrt{R}^l_{pred,.,t,k} - \sqrt{\lambda}^l_{.,t,k})^2$$

The CR time specific BPV ($BPV_t^{CR}$)

$$BPV_t^{CR} = \frac{\sum_l I(\sum_k FT^l_{pred,t,k} > \sum_k FT^l_{t,k})}{L}$$

and the CR global temporal $BPV$ ($BPV^{CR}_{Gt}$)

$$BPV^{CR}_{Gt} = \frac{\sum_l I(\sum_t \sum_k FT^l_{pred,t,k} > \sum_t \sum_k FT^l_{t,k})}{L}$$

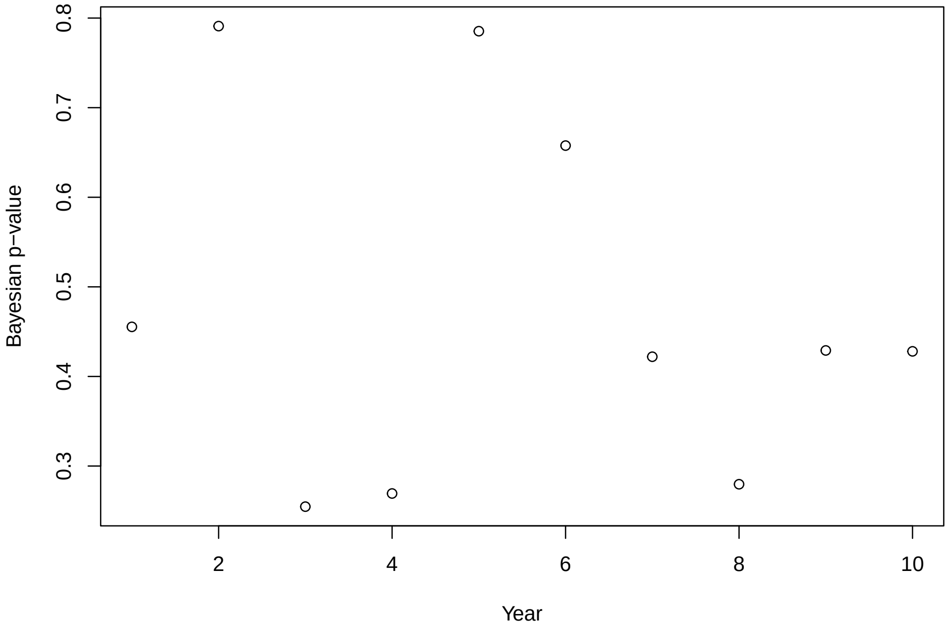


Figure S14: Year specific Bayesian p-value of resident individuals from the Monitoring Avian Productivity and Survivorship (MAPS) program.

CR global temporal known residence $BPV = 0.517$.

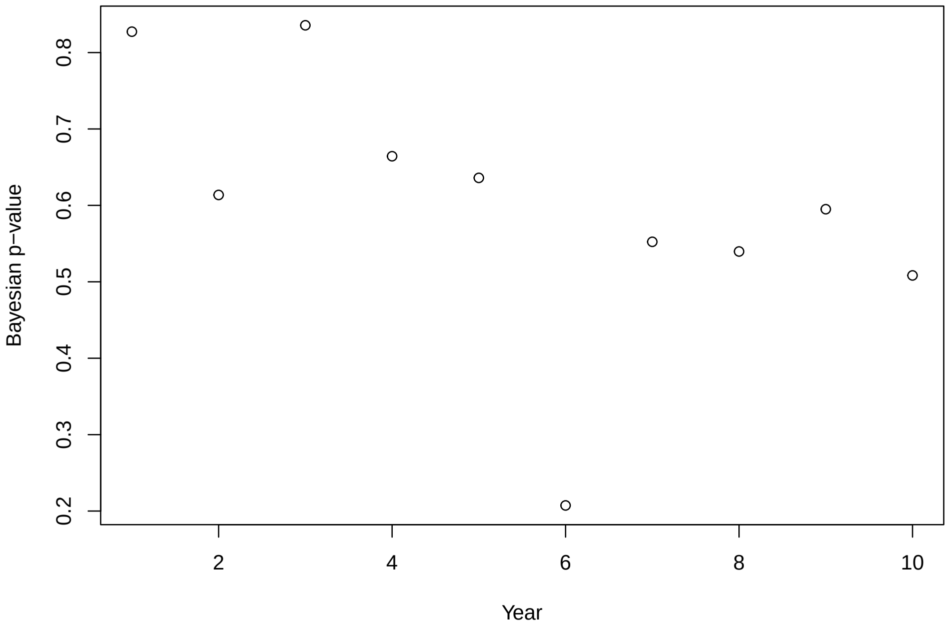


Figure S15: Year specific Bayesian p-value of possible transient and resident individuals from the Monitoring Avian Productivity and Survivorship (MAPS) program.

CR global temporal $BPV = 0.827$ for the mixture of transient and resident.

### References


Ahrestani, F. S., Saracco, J. F., Sauer, J. R., Pardieck, K. L., & Royle, J. A. (2017). An integrated population model for bird monitoring in North America. *Ecological Applications*, *27*(3), 916–924. https://doi.org/10.1002/eap.1493

Gelman, A., & Rubin, D. B. (1992). Inference from iterative simulation using multiple sequences. *Statistical science*, *7*(4), 457–472.

Pradel, R., Hines, J. E., Lebreton, J.-D., & Nichols, J. D. (1997). Capture-recapture survival models taking account of transients. *Biometrics*, 60–72.

Schaub, M., & Kéry, M. (2022). *Integrated population models: Theory and ecological applications with R and JAGS*. Academic Press.
